# A novel redox-active switch in Fructosamine-3-kinases expands the regulatory repertoire of the protein kinase superfamily

**DOI:** 10.1101/2020.01.13.904870

**Authors:** Safal Shrestha, Samiksha Katiyar, Carlos E. Sanz-Rodriguez, Nolan R. Kemppinen, Hyun W. Kim, Renuka Kadirvelraj, Charalampos Panagos, Neda Keyhaninejad, Maxwell Colonna, Pradeep Chopra, Dominic P. Byrne, Geert J. Boons, Esther van der Knaap, Patrick A. Eyers, Arthur S. Edison, Zachary A. Wood, Natarajan Kannan

## Abstract

Aberrant regulation of metabolic kinases by altered redox homeostasis is a major contributing factor in aging and disease such as diabetes. However, the biochemical mechanisms by which metabolic kinases are regulated under oxidative stress is poorly understood. In this study, we demonstrate that the catalytic activity of a conserved family of Fructosamine-3-kinases (FN3Ks), which are evolutionarily related to eukaryotic protein kinases (ePKs), are regulated by redox-active cysteines in the kinase domain. By solving the crystal structure of FN3K homolog from *Arabidopsis thaliana* (AtFN3K), we demonstrate that it forms an unexpected strand-exchange dimer in which the ATP binding P-loop and adjoining beta strands are swapped between two chains in the dimer. This dimeric configuration is characterized by strained inter-chain disulfide bonds that stabilize the P-loop in an extended conformation. Mutational analysis and solution studies confirm that the strained disulfides function as redox “switches” to reversibly regulate FN3K activity and dimerization. Consistently, we find that human FN3K (HsFN3K), which contains an equivalent P-loop Cys, is also redox-sensitive, whereas ancestral bacterial FN3K homologs, which lack a P-loop Cys, are not. Furthermore, CRISPR knockout of FN3K in human HepG2 cells results in significant upregulation of redox metabolites including glutathione. We propose that redox regulation evolved progressively in FN3Ks in response to changing cellular redox conditions. Our studies provide important new insights into the origin and evolution of redox regulation in the protein kinase superfamily and open new avenues for targeting HsFN3K in diabetic complications.

## Introduction

Glycation is a universal post-translational modification in which reducing sugars such as glucose and fructose are non-enzymatically added to free amine groups on peptides, proteins, and lipids. This non-enzymatic modification occurs endogenously in all living organisms, as well as exogenously in foods we consume (*1, 2*). Sugars attached to amine groups can undergo Amadori rearrangements to form stable linkages with biomolecules (*3–5*). Because such linkages can adversely affect biomolecular functions, organisms have evolved deglycation mechanisms (*6–8*) to repair the potentially toxic effects of reactive sugars. Fructosamine-3 kinases (FN3Ks) are a conserved family of deglycation enzymes that remove ribose and fructose sugars attached to surface-exposed lysine (Lys) residues (ketosamines) in proteins (*9–12*). They do this by catalyzing the transfer of the gamma phosphate from adenosine triphosphate (ATP) to the 3’ hydroxyl group in the ketosamine substrate. Phosphorylation of ketosamines by FN3Ks results in an unstable ketosamine 3-phosphate intermediate, which spontaneously decomposes into inorganic phosphate (*9*). Because the original unmodified Lys is regenerated as a consequence of FN3K deglycation activity, FN3Ks are believed to function as protein-repair enzymes (*6, 13*).

FN3Ks are conserved across the tree of life (*12, 14, 15*). Whereas simple eukaryotes and prokaryotes both contain a single copy of the FN3K gene, complex eukaryotes, including mammals, encode two copies, FN3K and FN3K-related protein (FN3KRP), presumably due to a gene duplication event in amphibians (*14*). Although the functions of FN3K homologs in lower eukaryotes and bacteria are yet to be equivocally established, it is proposed that they repair proteins modified by ribose-5-phosphate, a potent glycating agent generated by the metabolic pentose phosphate pathway (*6, 12, 14, 15*). FN3K activity is essential for normal cellular functions and uncontrolled activity can result in altered cellular homeostasis and disease (*16, 17*). For example, accumulation of 3-deoxyglucosone, a byproduct of human FN3K (HsFN3K) activity, is causatively associated with diabetic complications, such as retinopathy and neuropathy (*18, 19*) and the development of hepatocellular carcinoma (HCC) is dependent on the deglycation of nuclear transcription factor NRF2 by HsFN3K (*20*). Increased 3-deoxyglucosone levels also contributes to oxidative stress by inhibiting glutathione peroxidase, a potent cellular antioxidant (*21*). Thus, tight regulation of FN3K activity is crucial for maintaining cellular homeostasis.

We previously reported that FN3Ks belong to a large superfamily of protein kinase-like (PKL) enzymes that include eukaryotic protein kinases, small-molecule kinases, and atypical kinases (*22, 23*). FN3Ks are more closely related to small-molecule kinases, such as aminoglycoside kinase (APH) and choline kinases, than eukaryotic protein kinases (ePKs) and more distantly related to atypical pseudokinases such as Fam20C and SelO (*24–26*). Through quantitative comparisons of the evolutionary constraints acting on diverse PKL-fold enzymes, we demonstrated that ePKs share sequence and structural similarity with small-molecule kinases in the N-terminal ATP binding lobe, but diverge significantly in the C-terminal substrate-binding lobe (*22, 23, 27*). In particular, the extended activation segment connecting the ATP and substrate binding lobes that classically controls catalytic activity through conformational changes driven by reversible phosphorylation of serine, threonine and tyrosine residues (*28*) is unique to ePKs and absent in small molecule kinases, including FN3Ks (Fig. 1). A related paper from Byrne *et al*. demonstrates that in addition to reversible phosphorylation, oxidation and reduction of a conserved Cys residue in the activation segment is a much more common mode of Ser/Thr protein kinase regulation than had been previously appreciated (*29*).

**Fig. 1.**
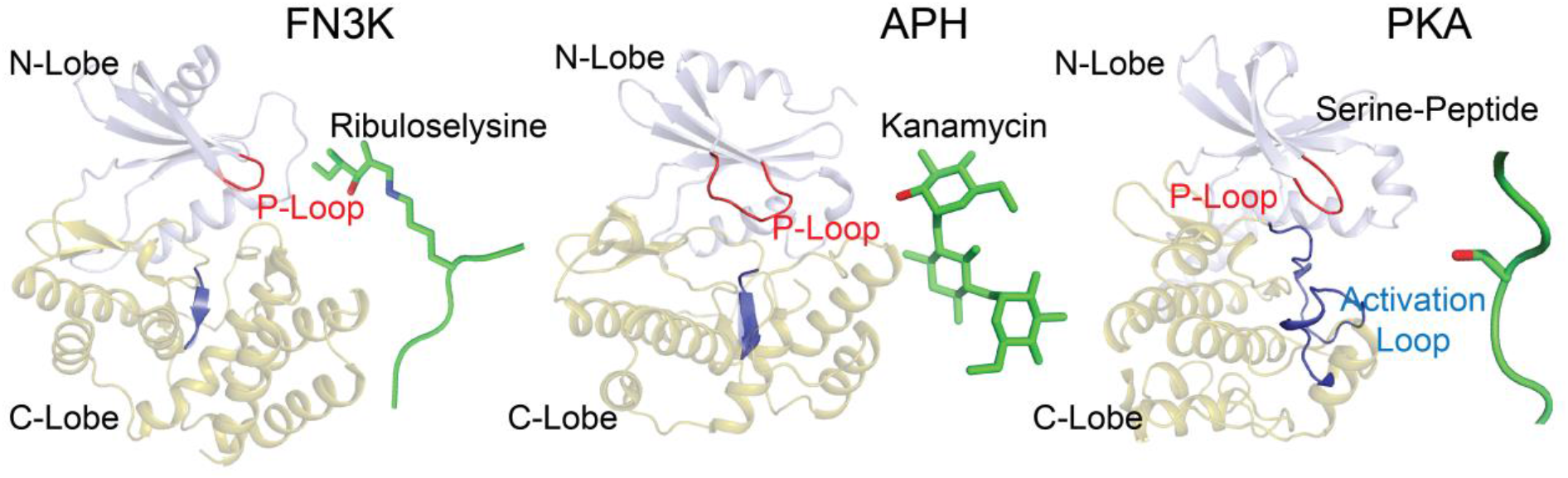
FN3K adopts a protein kinase fold. Comparison of the overall fold of FN3K from *T. fusca* (TfFN3K; PDB ID: 3F7W), aminoglycoside phosphotransferase (APH; PDB ID: 1L8T) (*85*) and protein kinase A (PKA; PDB ID: 1ATP) (*86*). The structures are shown as a cartoon where the N-lobe is colored in light blue and the C-lobe in olive green. The substrates are shown as either sticks (ribuloselysine, kanamycin) or cartoons (serine-peptide), and colored green. The oxygen atom on the hydroxyl group where the phosphate group is transferred is colored in red. The P-loop and activation loop are colored in red and blue, respectively.

Here, we uncovered a critical role for the ATP binding P-loop in the redox regulation of FN3Ks. By solving the crystal structure of a eukaryotic FN3K homolog, *Arabidopsis thaliana* FN3K (AtFN3K), we found that the P-loop is stabilized in an extended conformation by a Cys-mediated disulfide bond connecting two chains to form a covalently linked dimer in which the reduction of disulfides resulted in AtFN3K activation. Consistently, HsFN3K, in which the P-loop cysteine is conserved, is redox-regulated and displayed altered oligomerization when proliferating cells are exposed to acute oxidative stress.

We propose that redox control mediated by the P-loop Cys is an ancient mechanism of FN3K regulation that emerged progressively during FN3K evolution from bacteria to humans. Because many protein kinases contain an equivalent redox-active cysteine in the P-loop, our studies also have broad implications for understanding the structure, function, and evolution of all PKL-fold enzymes, in particular, tyrosine kinases, which contain a cysteine residue at the equivalent position in the P-loop. Our detailed mechanistic characterization of FN3Ks also opens new avenues for the design of FN3K-targeted small molecule inhibitors for diabetic complications associated with increased protein glycation.

## Results

### Crystal structure of AtFN3K reveals a previously unknown strand-exchange dimer

To investigate the structural basis for biological FN3K regulation, we solved the crystal structure of a plant FN3K homolog (AtFN3K) in complex with the ATP mimic adenylyl-imidodiphosphate (AMP-PNP) at a resolution of 2.37 Å using a multiple model molecular replacement strategy (table S1). The asymmetric unit of AtFN3K contains two molecules with a small degree of disorder at the N- and C-termini (residues 1-6 and 296, 297) (Fig. 2A). Each chain contains a well-ordered molecule of AMP-PNP in the active site, albeit with missing electron density for γ-phosphate (Fig. 2B). AMP-PNP is known to hydrolyze slowly over time, as shown previously for protein kinase A (*30*). Because the nitrogen atom in the β-phosphate of AMP-PNP could not be identified, we have modeled the ligand as adenosine diphosphate (ADP).

**Fig. 2.**
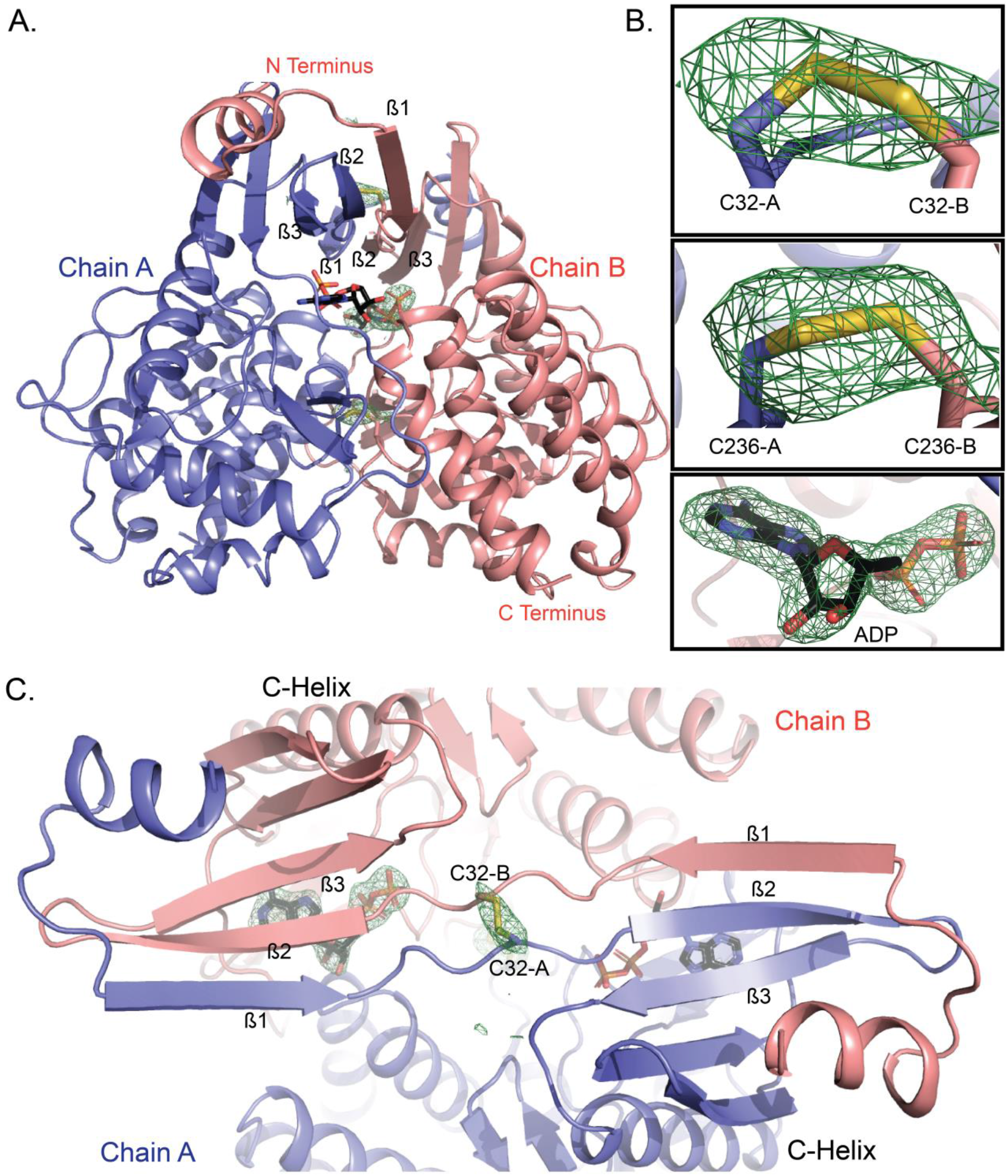
WT AtFN3K is a beta-strand exchange disulfide-mediated dimer. **(A)** Cartoon representation of the crystal structure of *A. thaliana* FN3K (AtFN3K) homodimer. The two disulfide bridges between two Cys^32^ and two Cys^236^ as well as the ADP molecules are shown as sticks. **(B)** Simulated annealing omit difference maps (*F_o_*-*F_c_*) calculated at 2.4 Å resolution and contoured at 4.5 rmsd. Maps were calculated after substituting both cysteines with alanine (top two panels) and removing the ADP molecule (bottom panel). **(C)** Top view of AtFN3K showing the beta-strand exchange.

AtFN3K adopts a canonical protein kinase fold (PKL-fold) with an N-terminal ATP binding lobe and C-terminal substrate-binding lobe. A comparison to the bacterial search model (PDB code 5IGS) shows that the chains superimpose 218 Cα atoms with an root mean square deviation (RMSD) of 3.3 Å (*31*). Due to the strong similarity between the two structures and the fact that the closely related aminoglycoside phosphotransferase structure from *E. coli* (5IGS) has already been described in detail (*32*), we focus here on the unique structural features of AtFN3K. Unlike other kinases, the AtFN3K structure is a “strand-exchange” dimer, in which the β1 strand in one chain forms an anti-parallel β-sheet with the β2 strand of the adjacent chain (Fig. 2C). In most PKL-fold crystal structures solved to date, the β1 strand forms an intra-chain anti-parallel beta-sheet with β2, and the P-loop connecting the two strands typically positions the ATP for catalysis, as observed in *Thermobifida fusca* (TfFN3K), aminoglycoside kinase and the prototypic ePK PKA (Figs. 1 and 3A). However, in the AtFN3K dimer, the P-loop is unfolded, and the extended P-loop conformation is stabilized by an intermolecular disulfide bond between Cys^32^ in chain A and Cys^32^ in chain B (Figs. 2B and 3A). The electron density of the Cys^32^ suggests that a small fraction of the disulfides may have been cleaved by X-rays during data collection, but we could not model both conformations with confidence given the relatively low 2.4 Å resolution. A second intermolecular disulfide is formed between Cys^236^ in each chain (Figs. 2B and 3B). Cys^236^ is located in the F-G loop, which is typically involved in substrate binding in both small-molecule kinases and protein kinases (Fig. 3B). In the AtFN3K dimer, the substrate-binding lobes are covalently tethered to create a unique interface, presumably for phosphorylating ketosamine and related substrates (Fig. 3C). The AtFN3K dimer buries nearly 2500 Å^2^ (16.4%) of solvent-accessible area of each monomer. The total solvation free energy (ΔG) gain upon dimerization is -36.2 kcal/mol, as determined by Protein Interfaces Surfaces and Assemblies (PISA) program (*33*), and is statistically significant(P-value of 0.014) with a total of 36 hydrogen bonds and two disulfide bridges at the dimer interface (fig. S1).

**Fig. 3.**
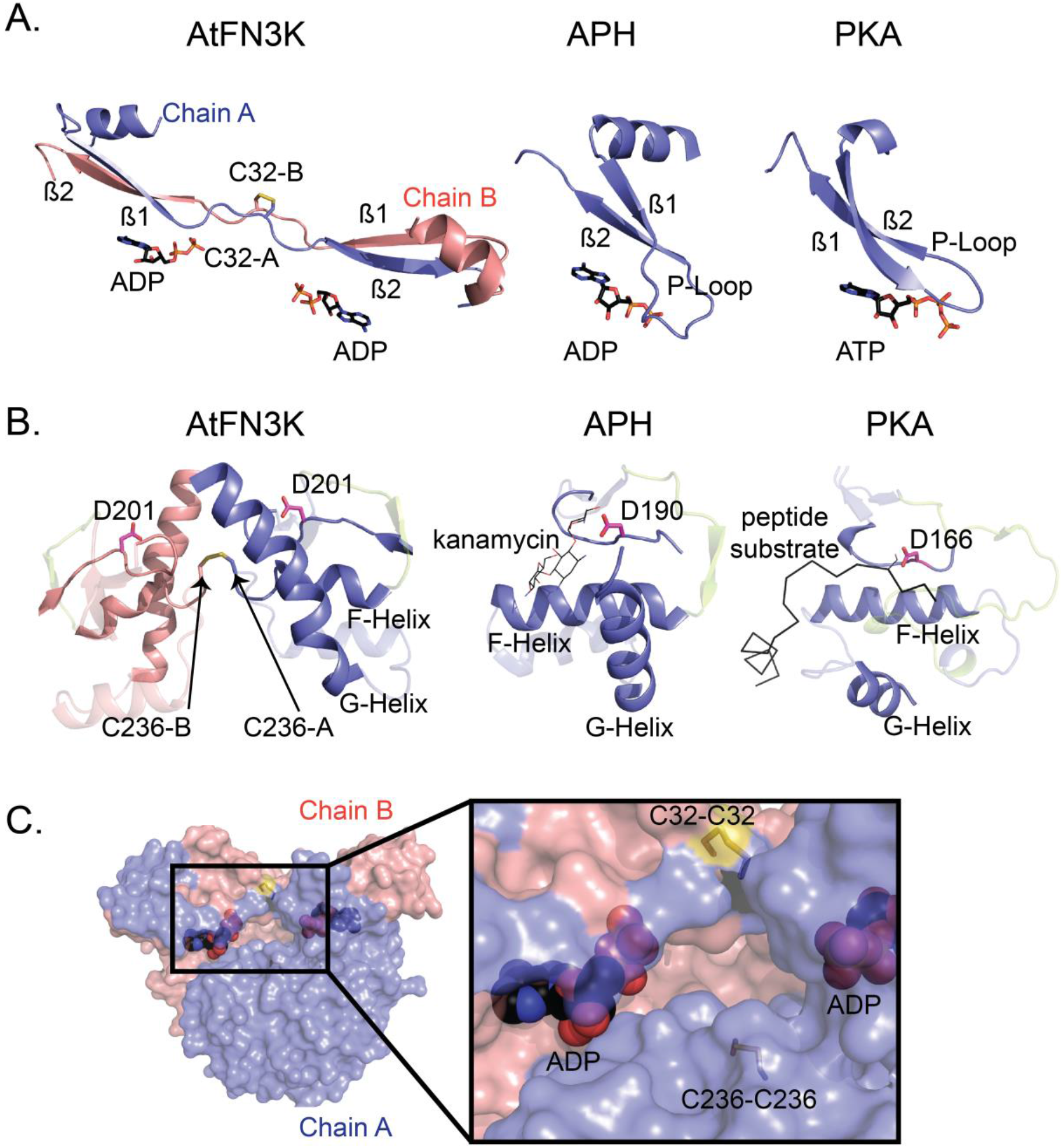
Comparison of the ATP- and substrate-binding regions in AtFN3K with APH and PKA. **(A)** Comparison of the P-loop of AtFN3K with that of APH and PKA. PDBs 1L8T and 1ATP were used for APH and PKA respectively. Carbon atoms of ADP and ATP molecules are colored in black, and the oxygen atoms are colored in red. Chain A and chain B of AtFN3K are colored in slate and salmon, respectively. **(B)** Comparison of the substrate binding lobe of AtFN3K with APH and PKA. Catalytic aspartate is shown as sticks with carbon atoms colored in magenta. The activation loop is colored in limon. The APH substrate kanamycin is shown as lines with carbon atoms colored in magenta. The PKA peptide substrate is shown as ribbonand colored in black. The serine residue in the peptide is modelled and shown as lines. PDBs used as in (A). **(C)** Surface representation of AtFN3K. Chains A and B and ADP-associated carbons are colored as described in (A). The two disulfide bridges, C32-C32 and C236-C236 are indicated with sticks with sulfur atoms colored in yellow.

### Disulfides in the AtFN3K dimer interface are conformationally strained

Disulfide bonds in proteins can be classified into various categories based on their geometry and torsional strain energy (*34–36*). Conformational analysis of intermolecular disulfides in AtFN3K indicates non-ideal geometry (Fig. 4A). In particular, the χ3 dihedral of -112.7° (Cβ- Sγ- Sγ- Cβ) for Cys^236^-Cys^236^ deviates from the peak distribution of ∼90° observed for disulfides in the Protein Data Bank (PDB) (Fig. 4B). Likewise, the Cα-Cα distance of 3.8 Å for Cys^32^-Cys^32^ is much shorter than 5.7 Å peak observed for Cα-Cα distances in the PDB (*37, 38*). These non-ideal geometries suggest that the intermolecular disulfides, particularly the Cys^236^-Cys^236^ disulfide, are conformationally strained.

**Fig. 4.**
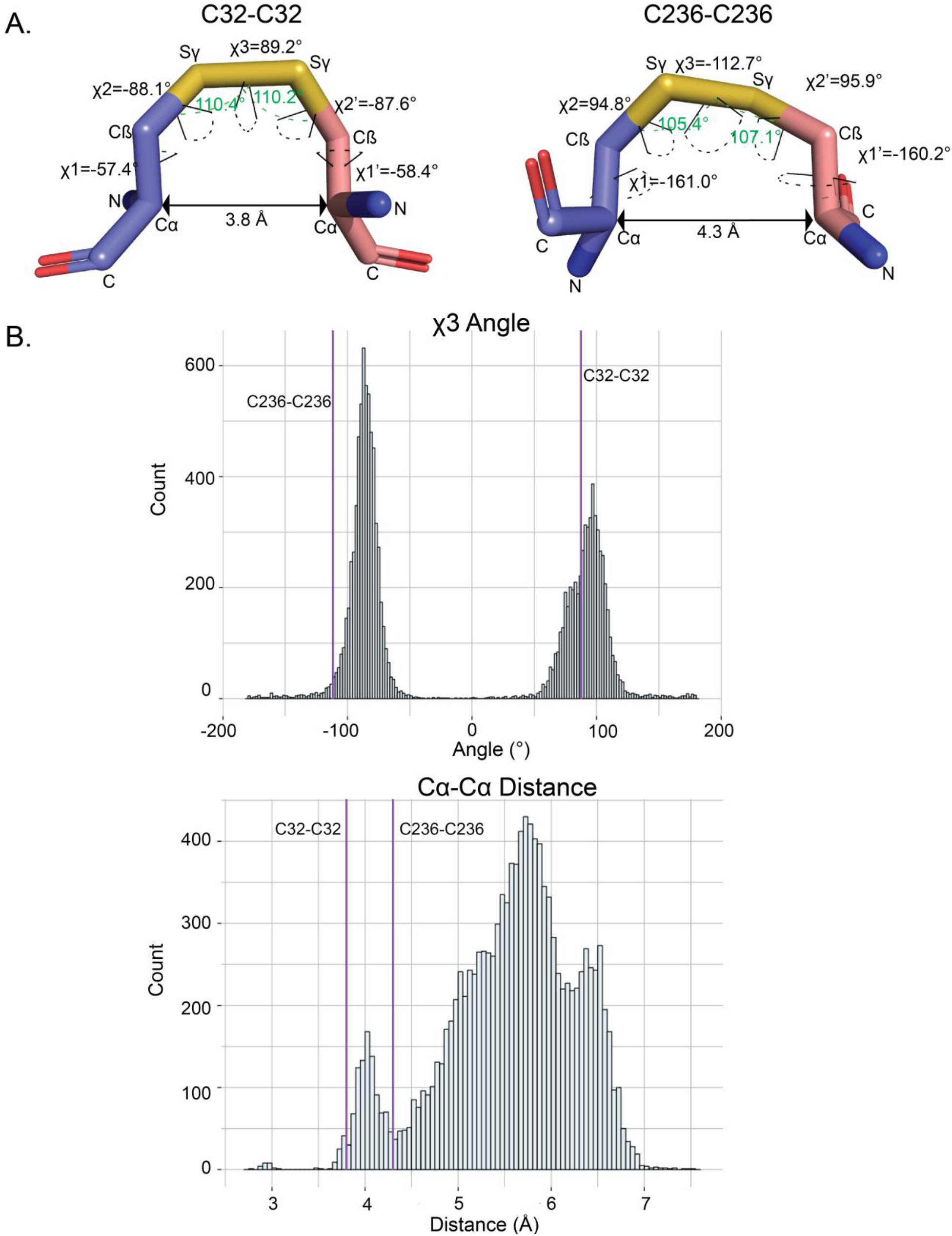
Geometric analysis of the disulfides in AtFN3K. **(A)** Distances and dihedral values labeled for the disulfide bridges between Cys^32^-Cys^32^ and Cys^236^-Cys^236^ between chain A and chain B. The angle (Cβ-Sγ-Sγ) is shown with a green dashed line. The dihedrals are shown with dashed line colored in black. PyMOL version 2.3 (*87*) was used to calculate the distances and the dihedral angles. **(B)** Distribution of the Cα-Cα distance and χ3 angle (Cβ-Sγ- Sγ- Cβ) for PDB structures with resolution less than 1.5 Å. The data was retrieved from (*65*). Values for the Cys^32^-Cys^32^ and Cys^236^-Cys^236^ disulfides in AtFN3K is represented by the purple vertical lines.

### P-loop cysteine is critical for the formation of disulfide-linked dimer species

Next, we wanted to test if either or both cysteines are essential for the observed disulfide-linked dimer. We mutated the cysteine residues individually and together to alanine residues and subjected them to non-reducing sodium dodecyl sulfate polyacrylamide gel electrophoresis (SDS-PAGE) analysis in the presence of redox agents, 1,4-dithiothreitol (DTT) and hydrogen peroxide (H_2_O_2_). The disulfide-linked dimer is absent in the P-loop cysteine mutants (C32A and C32A/C236A) but not in the C236A mutant (Fig. 5A), suggesting that the P-loop cysteine is critical for the observed disulfide-linked dimer. In addition, we identified multiple monomeric species on the gel, including a reduced monomer (M_Red_) and monomers with intramolecular disulfide (M_S-S_) (Fig. 5A). The presence of M_S-S_ band in the double cysteine mutant led us to hypothesize that the two remaining cysteines, Cys^196^ and Cys^222^ (Fig. 5B), in the kinase domain could form intramolecular disulfides. To test if an intramolecular disulfide is formed in solution or upon denaturation, we treated the size exclusion chromatography (SEC)-purified WT dimer with N-Ethylmaleimide (NEM) before denaturation. NEM blocks free thiols and prevents their oxidation and subsequent disulfide formation (*39*). The absence of M_S-S_ and the presence of M_Red_ in the NEM treated sample suggests that the M_S-S_ band is a consequence of denaturation (fig. S2). As a control, we also ran a non-reducing SDS-PAGE on the triple cysteine mutant (C32A/C236A/C196A), which is incapable of forming any intramolecular disulfides. As expected, only M_Red_ band was observed in the triple mutant (fig. S2), further suggesting that the M_S-S_ band is a consequence of denaturation during SDS-PAGE analysis. For this reason, we primarily focus on the inter-chain disulfides observed in the crystal structure.

**Fig. 5.**
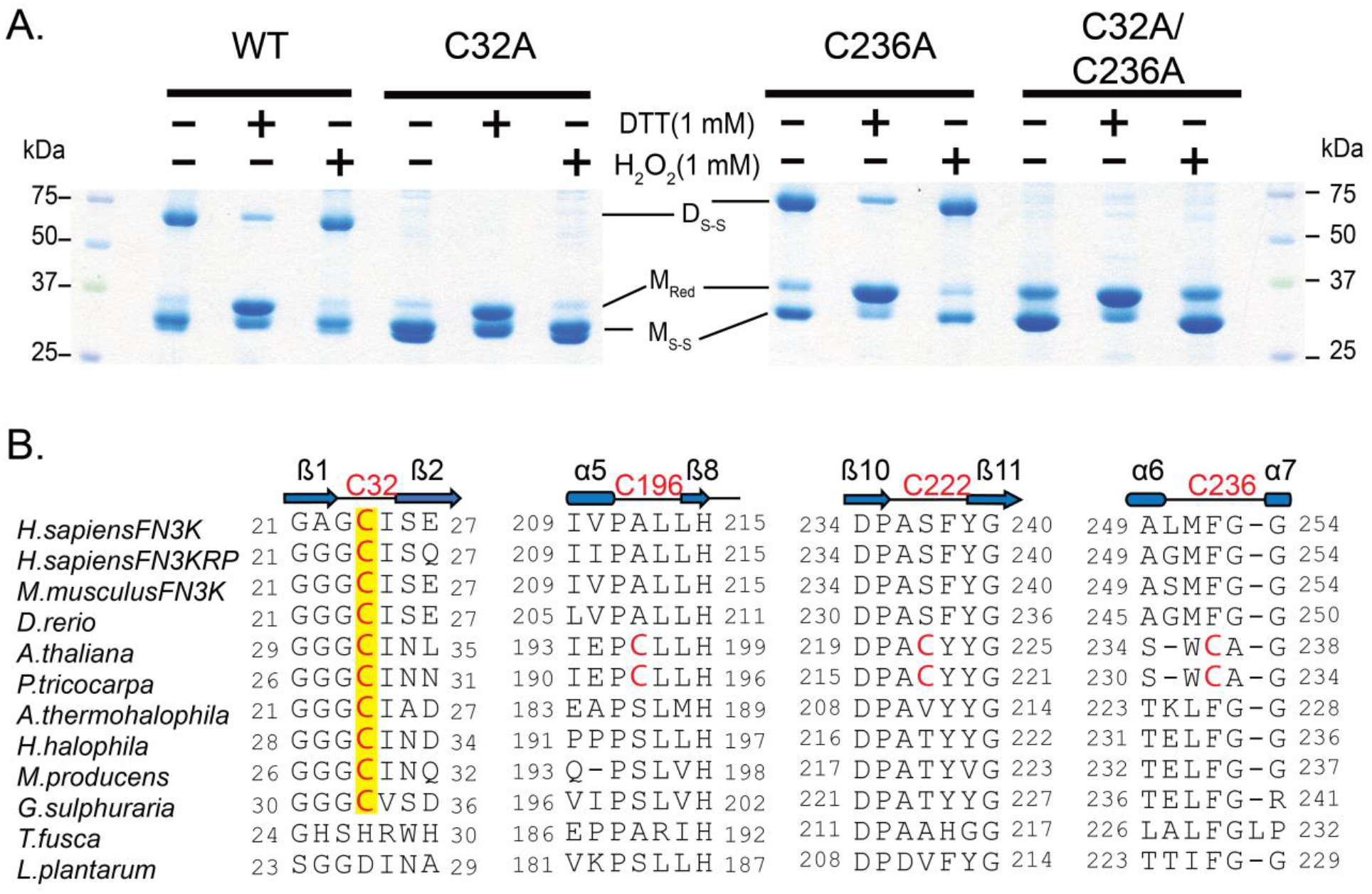
P-loop cysteine of AtFN3K (Cys^32^) is critical for the formation of disulfide-linked dimer species. **(A)** Non-reducing SDS-PAGE of WT and Cys-to-Ala mutant AtFN3K. 15 µg of protein was incubated with 1 mM DTT or 1mM H_2_O_2_ for 20 min and then subjected to SDS-PAGE under non-reducing conditions. D_S-S_: Disulfide linked dimer; M_Red_: Monomer Reduced; M_S-S_: Monomer with intramolecular disulfide. Blots are representative of 3 experiments. **(B)** Multiple sequence alignment of FN3K orthologs. Two additional cysteines (Cys^196^ and Cys^222^) specific to plant FN3Ks are shown. The alignment was generated using MUSCLE (*68*).

### AtFN3K WT dimer is activated by redox agents

In order to determine the functional state of the native enzyme (WT) in solution, we performed SEC on the purified protein. We identified two peaks, a dimer and a monomer (Fig. 6A and fig. S3A) with the dimer peak being dominant in solution. Compared to a reducing SDS-PAGE, the purified dimer resolved on non-reducing SDS-PAGE gel as two distinct bands (fig. S4, A and B). To test sensitivity to DTT, we performed a pyruvate kinase/lactate dehydrogenase (PK/LDH) assay on the dimer and monomer fractions in the presence or absence of 2 mM DTT. The activity of the dimer species increased nearly 40-fold in the presence of the reductant (Fig. 6B). In contrast, the monomer species was active but insensitive to DTT (Fig. 6B), suggesting that only the dimer species is redox-sensitive. The activity of the dimer species increased with increasing concentration of reduced glutathione (GSH), a physiological reductant (fig. S5A). We also performed PK/LDH assays on single and double cysteine to alanine or serine mutants. P-loop cysteine (C32A or C32S) mutants were less sensitive to DTT compared to C-lobe cysteine mutants (C236A/C236S) (fig. S5B). Moreover, AtFN3K WT also showed sensitivity to DTT when activity was measured independently using nuclear magnetic resonance (NMR)-based assay which measures ribuloselysine phosphorylation (fig. S5C).

**Fig. 6.**
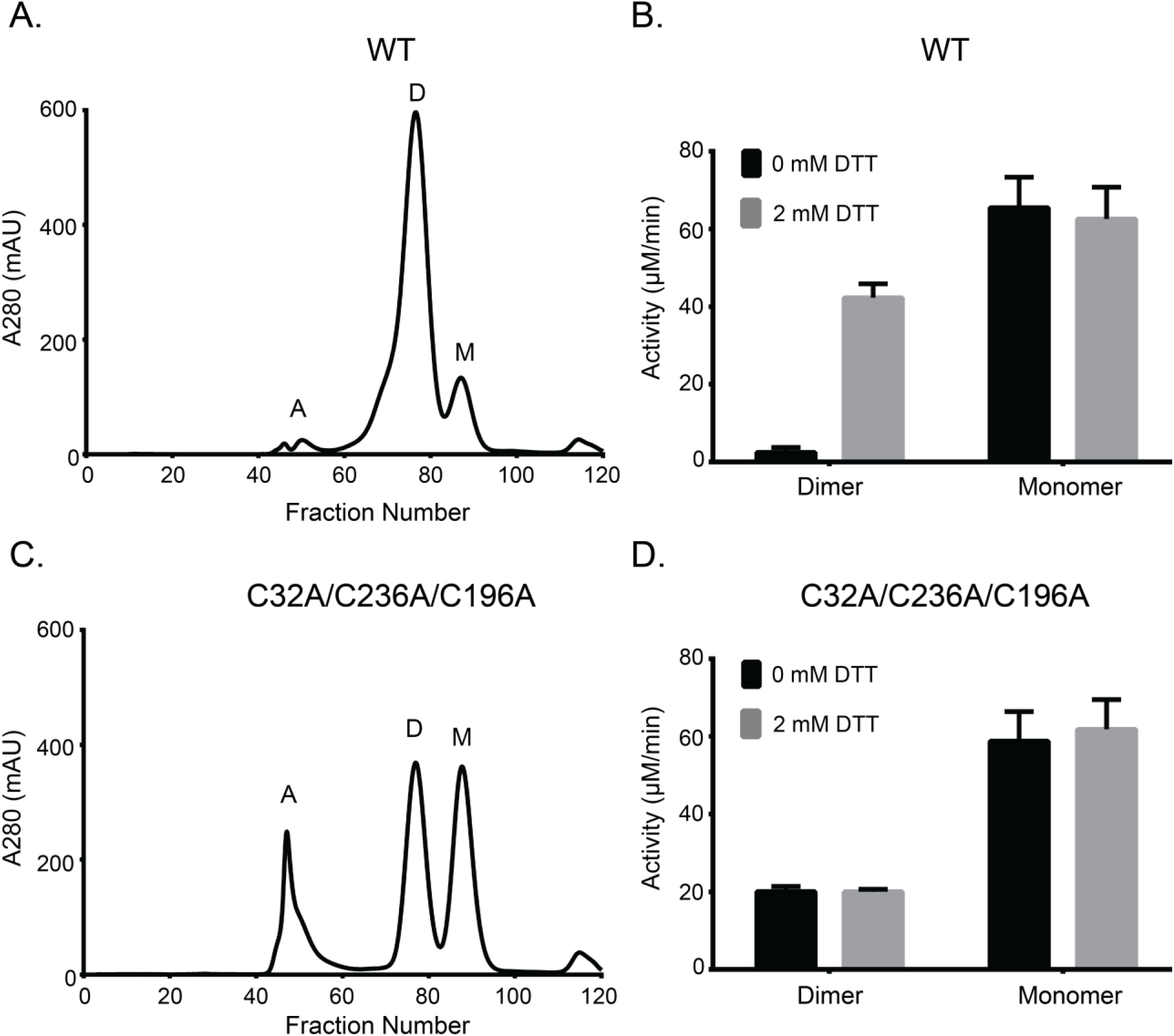
Both WT and triple-cysteine-mutant (C32A/C236A/C196A) AtFN3K exist as two distinct species in solution, and the WT dimer is redox sensitive. **(A)** Size exclusion chromatography (SEC) of AtFN3K WT protein. Each fraction was 1 ml in volume. A: aggregates, D: dimer, M: monomer. **(B)** PK/LDH assay using 1 µg of protein to assess the activity of WT protein in the presence or absence of 2 mM DTT. Ribulose-N-α-Ac-lysine was used as the substrate. Data are means ± standard error of six independent experiments. **(C and D)** As in (A) and (B), respectively, for the triple-cysteine-mutant protein.

To test if AtFN3K could dimerize without the cysteines, we performed SEC on the triple cysteine mutant (C32A/C236A/C196A). Like the native enzyme, we identified both dimer and monomer peaks (Fig. 6C). We hypothesized that the two species (monomer and dimer) would be insensitive to DTT since they are incapable of forming disulfide bridge. As expected, both the dimer and monomer species were insensitive to DTT (Fig. 6D). Together, these data suggest that AtFN3K can still dimerize in the absence of two cysteine residues (Cys^32^ and Cys^236^), which lie at the dimer interface.

To test whether the monomer and dimer species were in equilibrium, we first performed sedimentation velocity analysis on the purified C32A/C236A/C196A triple mutant. c(s) analysis revealed a distribution consisting of a monomer peak at 2.0 S and a dimer peak at 3.1 S which are predicted to be at 2.2 S and 3.6 S, respectively (fig. S3B). Glycerol is required to maintain the recombinant AtFN3K stably in solution and tends to form a self-gradient during centrifugation which will disproportionately reduce the S-values of sedimenting species. The monomer-dimer distribution is consistent with the size exclusion profile suggesting that the protein exists in two predominant species (Fig. 6C and fig. S3A). Secondly, we performed sedimentation velocity analysis across a moderate change (5x) in protein concentration but did not detect any changes in the ratio of monomers and dimers present (fig. S3B). To confirm that the species are not exchanging to any significant extent, we purified the dimer and monomer species using SEC and repeated the sedimentation velocity analysis (fig. S3C). The c(s) distribution of the purified dimer shows no significant dissociation but does contain a 1.3 S species that may correspond to a small amount of misfolded monomer. The c(s) distribution of the purified monomer suggests only a small amount of aggregation, as evidenced by a slight tailing of the c(s) peak toward larger s-values (fig. S3C).

### P-loop cysteine confers redox sensitivity in plant and mammalian FN3Ks

AtFN3K contains a chloroplast signal peptide N-terminus of the kinase domain, and removal of the signal peptide results in the localization of AtFN3K in different cellular compartments, including nucleus and mitochondria (fig. S8). Unlike AtFN3K, other orthologs only contain the kinase domain. Multiple sequence alignment of FN3K orthologs from diverse organisms indicates that while the P-loop cysteine (Cys^32^) is conserved across diverse organisms, the other three cysteines (Cys^236^, Cys^196^, and Cys^222^) are unique to plant FN3Ks (Fig. 5B). Notably, HsFN3K and human FN3K-related protein (HsFN3KRP) both contain an equivalent cysteine in the P-loop. We note that some protein tyrosine kinases that are distantly related to FN3Ks also conserve a Cys residue at the Cys^32^ equivalent position in the P-loop (fig. S9).

HsFN3K contains three additional cysteines, two of which are also present in HsFN3KRP. We expressed and purified HsFN3K and assayed its activity under reducing conditions. The activity of HsFN3K increased in the presence of DTT as observed for AtFN3K (Fig. 7A). Mutating the equivalent P-loop cysteine (Cys^24^) to alanine results in loss of sensitivity to DTT. We also tested the activity in the presence of GSH. The activity increased for the WT whereas no change was observed for the C24A mutant (fig. S6A). As a control, we expressed, purified and assayed AtFN3K homologs from *Thermobifida fusca* (TfFN3K) and *Lactobacillus plantarum* (LpFN3K), which contain a histidine and aspartate in place of the P-loop cysteine, respectively (Fig. 5B). Notably, TfFN3K and LpFN3K catalytic activity did not change in the presence of any of the redox agents tested (Fig. 7B). Also, non-reducing SDS-PAGE for the WT and C24A HsFN3K confirmed that the P-loop cysteine is essential for dimerization (fig. S6B).

**Fig. 7.**
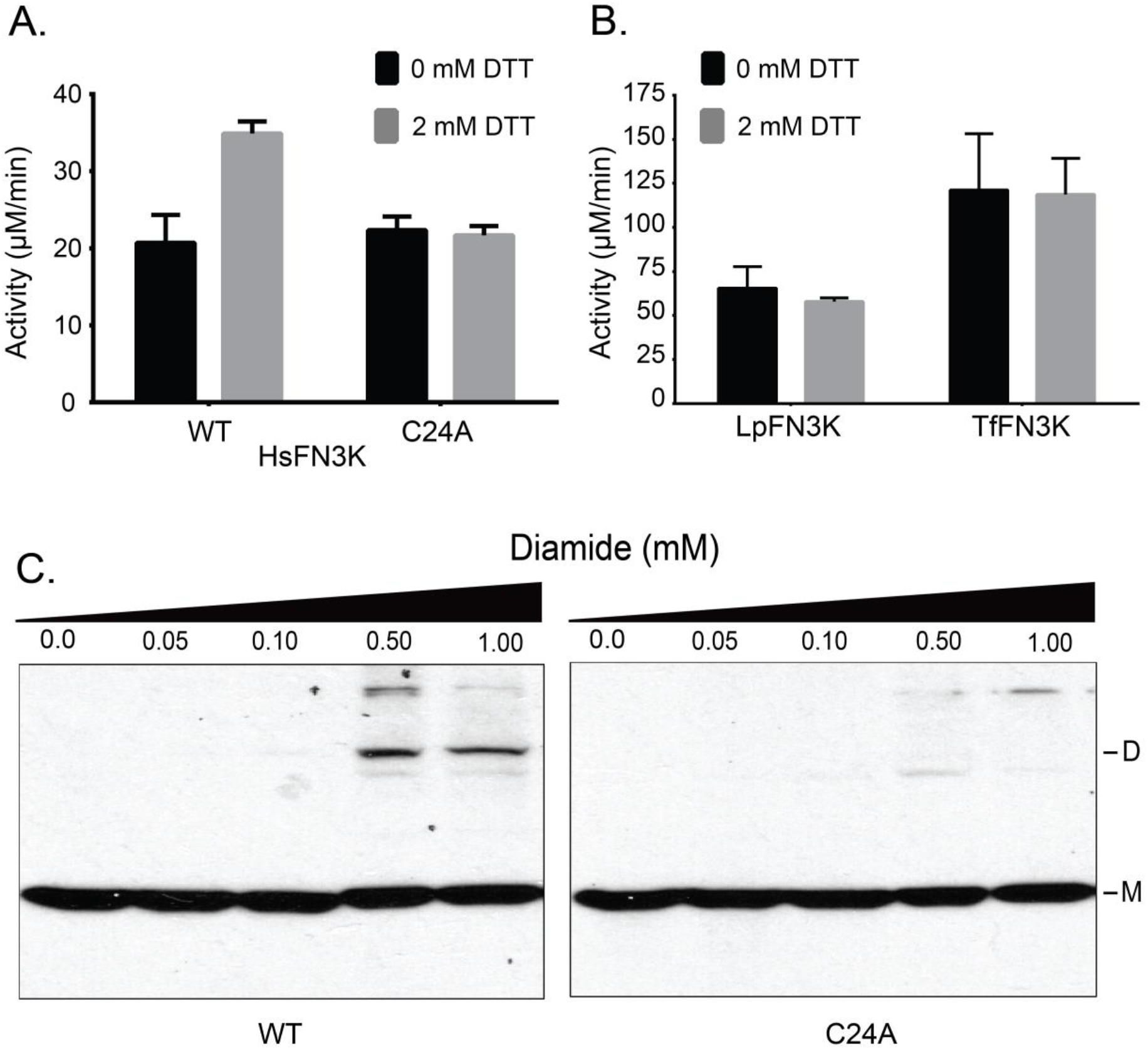
P-loop cysteine contributes to redox-sensitivity in HsFN3K. **(A and B)** PK/LDH assays performed with 10.0 μg of HsFN3K (A) or 1.0 μg each of TfFN3K and *L. plantarum* FN3K (LpFN3K) (B). Proteins were incubated with buffer (0 mM DTT) or 2 mM DTT, and ribulose-N-α-Ac-lysine was used as the substrate. Data are mean ± standard error of three independent experiments. **(C)** Effect of different diamide concentrations on transfected Flag-tagged WT and C24A HsFN3K in HEK293 cells. Total cell lysates were immunoblotted for Flag. D_S-S_: disulfide-linked dimer, M: monomer. Blot is representative of 3 experiments.

Finally, to extend these findings in mammalian cells, we expressed Flag-tagged HsFN3K in HEK293T cells using transient transfection. Cells were grown for 48 hours and treated with different concentrations of diamide. Diamide decreases the cellular concentration of GSH and promotes disulfide formation (*40*). It can also react with free thiols in proteins and lead to disulfide bond formation (*41*). Diamide treatment induced higher-order oligomerization of WT HsFN3K, but not for C24A mutant (Fig. 7C), suggesting that the P-loop cysteine confers redox sensitivity in both plant and mammalian FN3Ks.

### FN3K CRISPR knockout alters redox-sensitive cellular metabolites

Cellular functions of FN3Ks are currently unknown. Our analysis of FN3K expression in various cancer cell lines identified significant overexpression in the liver and eye cancer cells (fig. S11). To assess the functional significance of this increased expression, we generated a Clustered Regularly Interspaced Short Palindromic Repeats (CRISPR) knockout of HsFN3K (FN3K-KO) in HepG2 liver cancer cell line (fig. S7) and compared the metabolome of WT and FN3K-KO cells using untargeted ^1^H nuclear magnetic resonance (^1^H NMR) metabolomics. This revealed a significant difference in metabolite abundance in FN3K-KO compared to WT (Fig. 8, A and B, table S2). Of special note, glutathione and lactate levels were increased in FN3K-KO cells relative to wild-type HepG2 cells, while pantothenate, phosphocreatine/creatine ratio, aspartate, glycine, and serine levels were decreased. Glutathione is a major cellular redox regulator (*42*), while intracellular levels of pantothenate, glycine, serine, and aspartate, are known to be reactive to the redox status of the cell (*43–45*). Additionally, phosphocreatine/creatine ratio and lactate control ATP production and glycolysis, respectively (*46*). The enrichment of these metabolites suggests potential links between FN3Ks, redox levels, and ATP production.

**Fig. 8.**
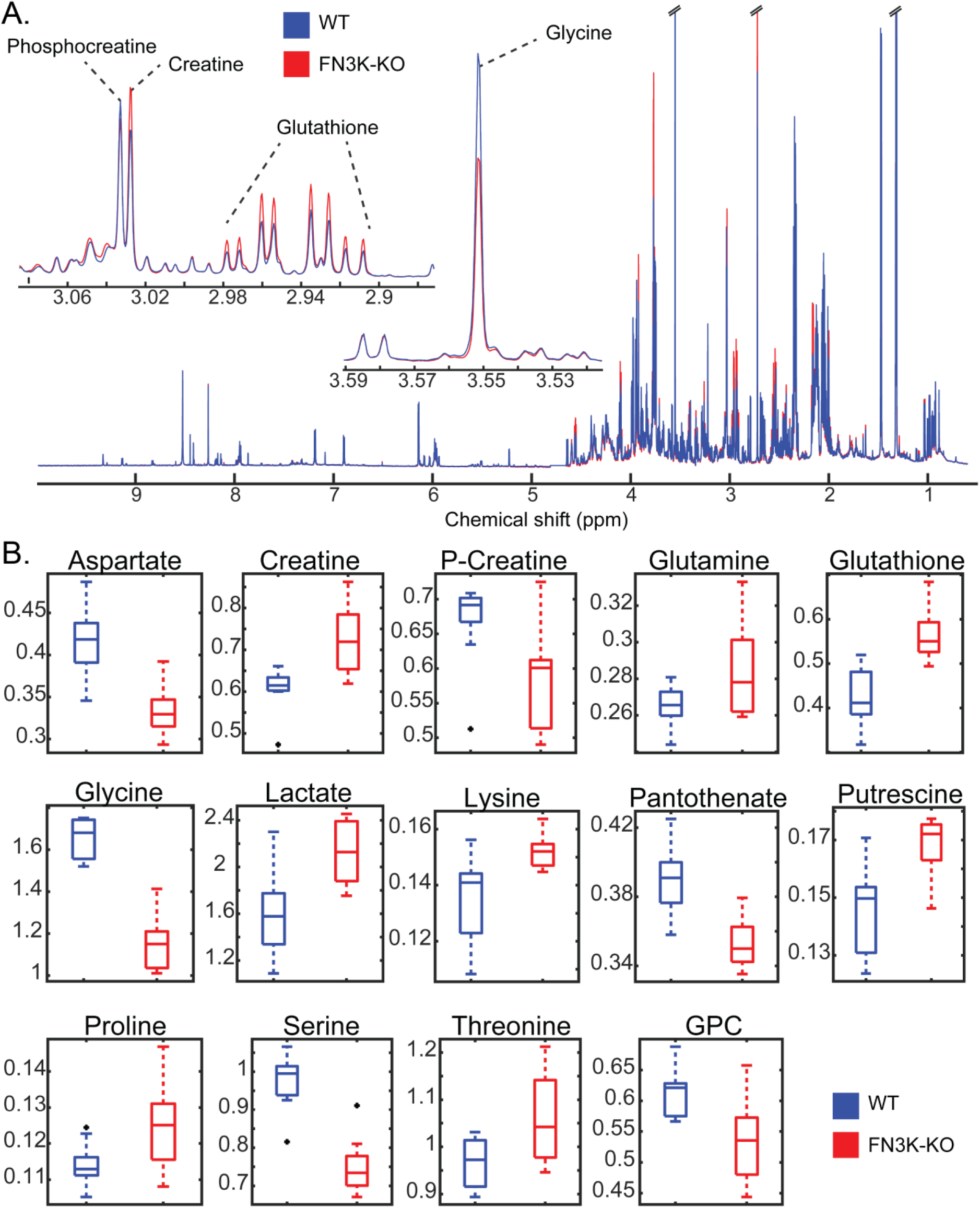
Redox-sensitive metabolites are altered in HsFN3K knockout cells. **(A)** ^1^H NMR spectra of WT and FN3K-knockout (FN3K-KO) HepG2 cells. Traces are the average for each group (WT N=10, FN3K-KO N=9). Insets highlight examples of regions containing annotated metabolites observed to be significantly different between cell lines. **(B)** Box and whisker plots of significant (FDR p-value < 0.05) metabolites annotated with highest confidence. Black points indicate outliers. Two-tailed T-test was performed, and the p-values were corrected for false discovery rate using the Benjamini-Hochberg method.

## Discussion

Here we report the structural basis for redox regulation in an ancient family of phosphorylation-based enzymes associated with protein repair. The crystal structure of *Arabidopsis thaliana* FN3K revealed a novel strand exchange dimer in which two chains are covalently linked by disulfide bonds emanating from the P-loop in the ATP binding lobe and F-G loop in the substrate binding lobe. We propose that these disulfide bridges are conformationally strained and are essential for maintaining AtFN3K in an inhibited dimeric conformation. Our studies also show that inhibited native AtFN3K (WT dimer) is activated more than 40-fold under reducing conditions. We speculate that reduction of the disulfides releases the constraint on the P-loop and the substrate-binding F-G loop to allow ATP and substrate access, and catalysis in the dimeric state. Thus, AtFN3K toggles between an inhibited “oxidized” dimer state and active “reduced” dimer to phosphorylate ketosamine substrates. Using SEC, we also identified a monomeric species insensitive to DTT. Although crystallization attempts to isolate the monomeric species have been unsuccessful, we predict that a monomer would adopt a canonical kinase fold without any beta-strand exchange. As a result, both P-loop and substrate-binding loops cannot be constrained through disulfide bridges, preventing redox regulation.

Similarly, SEC applied to the C32A/C236A/C196A mutant revealed the presence of both dimeric and monomeric species, indicating that the enzyme can dimerize without these cysteines. PISA analysis of the crystal structure shows that nearly 2500 Å^2^ (16.4%) of each monomer is buried as part of the dimer interface. Sedimentation velocity analysis further revealed that the two species were distinct and do not exist in equilibrium.

Our cellular studies show that dimerization and higher-order oligomerization of HsFN3K is altered after exposure to the thiol-oxidizing agent, diamide. Consistently, an unbiased proteomic study identified a sulfenylated form of the P-loop cysteine in HsFN3KRP in HeLa cells (*47*). Moreover, a recent proteomic study identified reversible oxidative modifications of the P-loop cysteine in FN3K and FN3KRP in both mouse and human tissues (*48*), suggesting that a regulatory mechanism of reversible P-loop oxidation is likely operative in cells and evolutionarily conserved.

The identification of redox-active cysteines in FN3Ks also opens up the possibility of a feedback regulation mechanism in which FN3K activity is controlled by its catalytic byproduct, 3-deoxyglucosone, which is known to contribute to oxidative stress (*21*). We speculate that the accumulation of advanced glycation end products such as 3-deoxyglucosone drives the conformational ensemble of FN3K species towards an inactive dimeric form through the oxidation of the P-loop cysteine, whereas reduced levels maintain FN3K in an active reduced form (Fig. 9). Such a feedback inhibition mechanism might explain how the essential deglycation functions of FN3K are carried out in cells without contributing to oxidative stress. Consistent with this view, our biochemical and mutational studies confirmed that inhibited dimeric species predominate in the absence of reducing agents, while active dimeric or monomeric species may dominate in the presence of reducing agents. Moreover, our comparative metabolomics study of the HsFN3K WT and KO in HepG2 liver cells revealed alteration in redox-sensitive metabolites, namely glutathione. Increased cellular glutathione in the KO relative to WT suggests potential links between FN3K functions and oxidative stress response that needs to be tested in future studies. In particular, delineating the pathways that link the redox active metabolites identified in our study with HsFN3K mediated deglycation functions reported recently in oncogenesis (*49*) will be a major goal moving forward.

**Fig. 9.**
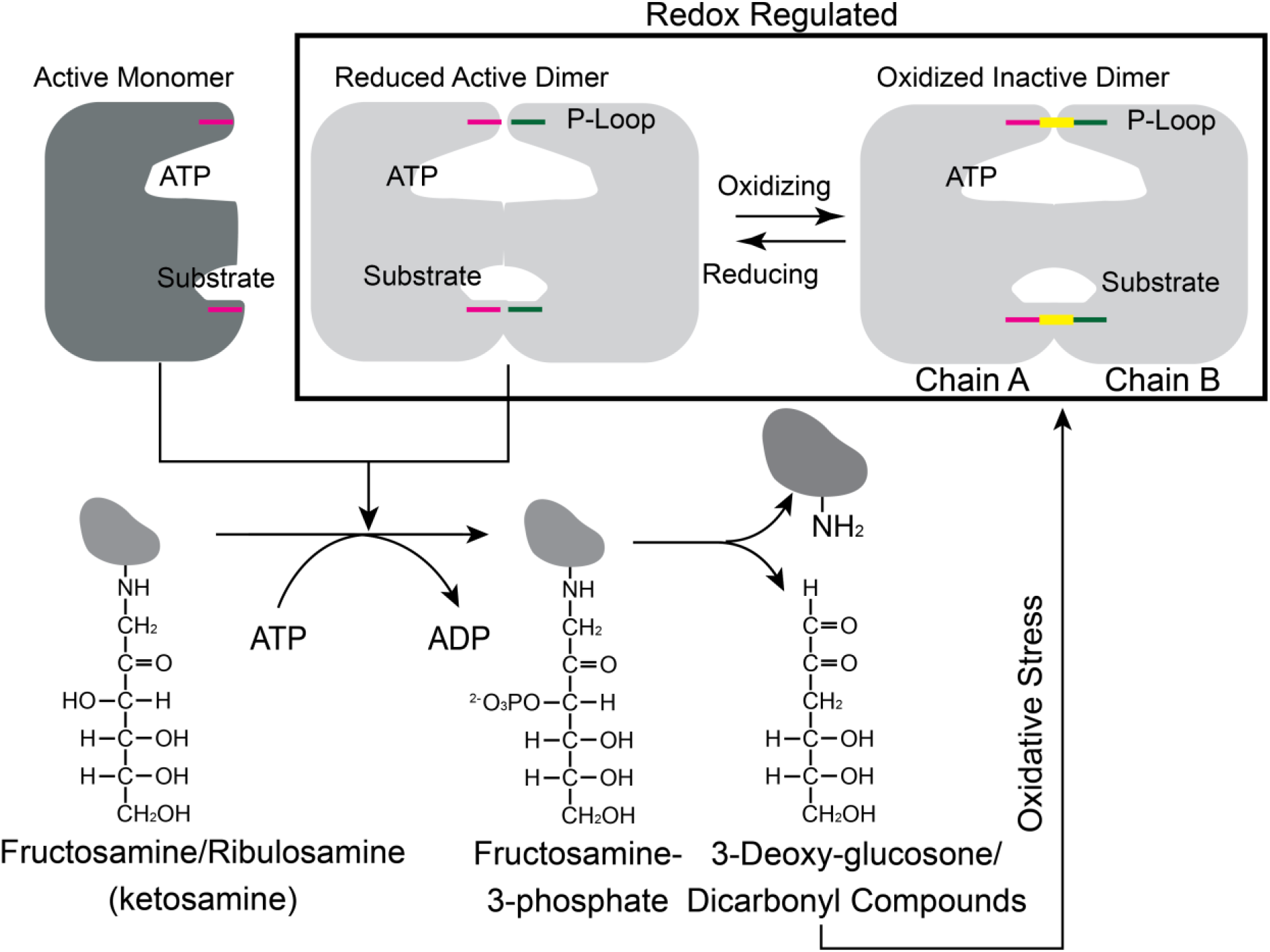
Proposed redox feedback regulation of plant and mammalian FN3Ks. Cartoon showing the possible relationship between redox regulation of FN3K activity and its physiological function. The disulfide is colored in yellow.

Redox control of FN3K may also explain the altered glycated patterns observed in plant and human proteomes (*50–53*) and facilitate deglycation functions to be carried out in different cellular compartments. Our studies on AtFN3K (without the signal peptide) revealed nuclear and mitochondrial localization (fig. S8). HsFN3K is localized in the mitochondria (*54*) and has been detected in red blood cells and serum samples. Mitochondria generates reactive oxygen species (ROS), which can mediate redox signaling through oxidation of SH groups in cysteine residues in proteins (*55–57*). The presence of a redox-sensitive P-loop cysteine (Cys^24^) in HsFN3K further suggests that redox regulation of FN3K is physiologically relevant.

The P-loop Cys is conserved amongst FN3K orthologs from diverse prokaryotic and eukaryotic organisms, and our studies indicate that the FN3K homolog in humans (HsFN3K) is also redox-sensitive. In contrast to the activation loop, which appears to have evolved as a flexible motif for eukaryotic protein kinase regulation by phosphorylation, the P-loop is fundamentally conserved in diverse ATP fold enzymes, where it plays a critical role in clamping ATP and positioning it for efficient catalysis. Here we show that conformational control of the P-loop by reactive cysteines is an ancient mode of regulation. Notably, several eukaryotic tyrosine kinases such as SRC, FGFR, YES1, and FYN conserve a cysteine residues at the Cys^32^ of AtFN3K equivalent position in the P-loop (fig. S9), suggesting potential redox regulation of these kinases as well. Indeed, both SRC and FGFR have previously been shown to be redox-regulated through Cys oxidation mechanisms in the P-loop (*58, 59*). However, whether these kinases also form a strand exchange dimer like AtFN3K is not known, and it will be interesting to attempt to solve the structure of Tyr kinases crystallized under different oxidizing conditions.

Finally, HsFN3K and HsFN3KRP are implicated in diseases such as diabetic neuropathy and retinopathy, and more recently in cancer (*49*). However, developing therapeutic strategies for FN3Ks is potentially a double-edged sword because inhibition of FN3K activity can result in the accumulation of glycated proteins, while activation of FN3Ks might result in the accumulation of 3-deoxyglucosone, a potent generator of oxidative stress. Based on our findings, we propose a combinatorial drug development approach, in which the active monomeric species and the reduced dimeric species are targeted through the development of thiol-interacting agents. These might be developed to help normalize cellular protein glycan homeostasis, depending upon the indication and redox balance in the appropriate tissue and the availability of redox-active Cys residues in FN3K targets.

## Materials and Methods

### Expression and purification of AtFN3K WT and Cys-to-Ala mutants

AtFN3K inserted into pET-15b expression vector was provided by Prof. Emile Van Shaftingen (Université Catholique of Louvain, Brussels, Belgium). Site-directed mutagenesis was performed using Q5 Hot Start High Fidelity Kit from New England Biolabs (NEB). Vectors were transformed into E. coli BL21(DE3) pLysS (Roche) cells and co-expression with GroEL was performed at 16°C for 18 hours in 500 ml of Luria–Bertani medium containing 100 μg/l ampicillin and 25 μg/l chloramphenicol. The bacterial extract was resuspended in 50 mM sodium phosphate buffer, pH 7.8, 300 mM NaCl, 10 % v/v glycerol (buffer A) with 1mM Phenylmethylsulfonyl Fluoride (PMSF) and 3 mM 2-mercaptoethanol (βME). The solution was sonicated on ice and the lysate was centrifugated at 15,000 x g for 45 minutes at 4°C. The supernatant was then passed through 0.5 ml of Talon® resin previously equilibrated with buffer A without βME. The column was washed with 20 ml of buffer B (50 mM Sodium phosphate buffer, pH 7.8, 1 M NaCl,10 % v/v glycerol) followed by a wash with 100 ml of buffer B containing 10 mM imidazole. The protein was then eluted with buffer A without βME containing 250 mM imidazole. The purity of the protein was checked by SDS-PAGE, and the fractions with the protein were dialyzed against buffer A without glycerol and later against buffer C (25 mM HEPES, pH 7.8, 300 mM NaCl, 10% v/v glycerol). The protein was snap-frozen with liquid nitrogen and stored at -80°C. Recombinant His-HsFN3K was co-expressed with GroEL in *E. coli* as well.

### Crystallization, phasing and refinement

Recombinant AtFN3K WT was dialyzed into a buffer containing 50 mM NaCl and 15 mM HEPES at pH 8.0, concentrated to ∼30 mg/mL and quantified using the extinction coefficient (Abs 0.1%: 1.576) calculated by Protparam (*60*). The protein was crystallized at 20°C using hanging drop vapor diffusion with 2 μl drops (1:1 protein to reservoir ratio). Crystals grew from a reservoir of 2 M ammonium sulfate, 0.1 M MES pH 5.5, 3.28 mM MgCl_2_ and 1.9 mM AMP-PNP. Crystals were cryoprotected with the reservoir solution supplemented with 10 % cryoprotectant (1:1:1 ratio of ethylene glycol, DMSO, and glycerol) and flash cooled in liquid nitrogen. The diffraction data were collected at the SER-CAT 22-ID beamline at the Argonne National Laboratory using a Rayonix 300HS detector and processed using XDS (*61*). Five percent of the data was set aside for cross validation. The structure was solved by molecular replacement in Phenix (*62*) using a multiple model template (structures from *T. fusca* (PDB code 3F7W), *H. somnus* (PDB code 3JR1) and *E. coli* (PDB code 5IGS) were used) with trimmed side chains and weighted according to structure and homology. Sequence matching and model building was performed using Phenix Autobuild (*62*). Automated refinement in Phenix (*62*) and iterative manual fitting using Coot (*63*) produced the final model (table S1). The B-factors were refined using TLS (*64*).

### Distribution of χ3 angle and Cα-Cα distance

We downloaded the dataset of culled crystal structures with unique disulfides from (*65*) and selected structures with a resolution less than or equal to 1.5Å to plot the histogram in Fig. 4B. *SDS-PAGE*

15 μg of the enzyme was incubated with distilled H_2_O or redox agents (DTT/H_2_O_2_) in a total volume of 15 μL. After 20 min, 5 μL of 4X Sample Buffer without βME (Beta Mercaptoethanol) was added and the protein was denatured at 100°C for 5 minutes. 10 μL of the sample was loaded onto a 12% SDS-PAGE gel. The gels were stained with Coomassie for an hour and de-stained with water for several hours. Similarly, 3-5 μg of the enzyme (15 μL volume) was incubated with 20 mM NEM or buffer (15mM HEPES, pH 7.8, 300 mM NaCl, 10 % v/v Glycerol) for 5 mins at room temperature before adding 4X Sample Buffer with (reducing) or without (non-reducing) 2.8 M βME.

### Sequence Alignment

Representative FN3K orthologs were downloaded from Uniprot (*66*), National Center for Biotechnology Information (NCBI) (*67*) and aligned using MUltiple Sequence Comparison by Log-Expectation (MUSCLE) (*68*). Eukaryotic protein kinases (ePKs) with P-loop cysteine conserved were identified using previously curated alignment profiles that included ePKs and small-molecule kinase sequences, including FN3Ks (*22, 23, 69, 70*).

### Expression analysis

Human tumor expression data was obtained from the National Cancer Institute’s Genomic Data Commons (GDC) Data Portal (downloaded Feb 10, 2017) (*71*). RNA-seq data was available for 11,574 samples (10,833 tumor samples and 741 normal samples). Read count for each gene was normalized by the Fragments Per Kilobase of transcript per Million mapped reads upper quartile (FPKM-UQ) method (*72*). Z-score for each gene was calculated based on deviation from the normalized mean expression levels (in logarithmic scale) for each sample.

### Size exclusion chromatography (SEC)

Approximately 4.5 mg of AtFN3K WT and triple cysteine mutants purified using co-expression with GroEL/ES chaperone (9 mg/ml in concentration) was passed through HiLoad 16/1600 Superdex 200 pg column at a flow rate of 1 ml/min with fractionation volume of 1 ml. 25 mM HEPES, pH 7.8, 300 mM NaCl, 10 % v/v glycerol was used as running buffer. SEC was performed at 4 °C. For BSA-lysozyme standard, 250 µl of 4 mg/ml of BSA and lysozyme each in running buffer were mixed and loaded on the column. For AtFN3K WT and triple cysteine mutants purified without GroEL chaperone (fig. S3A), SEC was performed at a flowrate of 0.15 ml/min.

### PK/LDH enzyme assays

1.0 μg (5 μL) of FN3K homologs (At, Tf, Lp)/mutants/dimer and 10 μg (5μL) of HsFN3K, 28 mM of ribulose-N-α-Ac-lysine (10 μL) in the presence or absence of 20 mM DTT (5 μL; final concentration: 2 mM) were mixed with 20 μl of solution prepared by mixing the following: 150 µl of 5X Kinase Buffer (160 mM HEPES, pH 7.4, 80 mM MgCl2, 1.2 M NaCl, 40 % v/v Glycerol), 15 µl of 250 mM phosphoenolpyruvic acid (PEP), 45 µl of PK/LDH mix [600-100 units/ml pyruvate kinase (PK), 900-1400 units/ml lactic dehydrogenase (LDH)], 90 µl of 37.5 mM nicotinamide adenine dinucleotide (NADH). The reaction was started by adding 10 μL of 5 mM ATP (final: 1mM). The final reaction volume was 50 μL per well. The 96well plate was immediately placed in a plate reader (Biotek Synergy H4) and the absorbance measured at 340 nm at 35°C continuously for two and half hours. Proteins were stored in Buffer D (25 mM HEPES, pH 7.4, 300 mM NaCl, 10% v/v glycerol). Buffer D was also used as mock buffer as needed.

### NMR real time assay

100 μg of protein was incubated with 2 mM of ribuloselysine and 600 μM of ATP in 25 mM HEPES, pH 8.0, 5 mM of MgCl_2_ prepared in D_2_O. The reaction was monitored for 50 minutes by acquiring subsequent 1D 1H PURGE (Presaturation Utilizing Relaxation Gradients and Echoes) spectra. All spectra were acquired at 22 °C on an 800 MHz Avance Neo (Bruker) NMR spectrometer equipped with a z-gradient triple resonance TCI cryoprobe and normalized and references using ^1^H signals of 4,4-dimethyl-4-silapentane-1-sulfonic acid (DSS). 32 scans were acquired during each experiment with an acquisition time of 1.25 sec. The spectra were processed using NMRpipe and Matrix Laboratory (MATLAB).

### Sedimentation velocity

The triple cysteine mutant (C32A/C236A/C196A) was dialyzed into 25 mM HEPES (pH 7.8), 300 mM NaCl, 2 mM MgCl_2_, 100 μM ATP, and 1 M glycerol, and the protein was quantified using an Agilent 8453 UV/vis with an ε280 of 55810 M-1cm-1 determined by ProtParam (*60*). The sample was diluted to a final protein concentration of 10 μM then loaded into a 12 mm double-sector Epon centerpieces equipped with quartz windows. The cell was loaded into An60 Ti rotor and equilibrated for 1 hour at 20°C. Sedimentation velocity data were collected using an Optima XLA analytical ultracentrifuge (Beckman Coulter) at a rotor speed of 50000 RPM at 20°C. Data were recorded at 280 nm in radial step sizes of 0.003 cm. SEDNTERP (*73*) was used to model the partial specific volume of AtFN3K (0.713293 mL/g), as well the density (1.0353 g/mL) and the viscosity (0.013277 P) of the buffer. SEDFIT (*74*) was used to analyze the raw sedimentation data. Data were modeled as continuous c(s) distribution and were fit using baseline, meniscus, frictional coefficient, and systematic time-invariant and radial-invariant noise. Fit data for the experiment had an RMSD in the range of 0.005–0.007AU. Predicted sedimentation coefficient (s) values for AtFN3K monomer (2.2 S) and dimer (3.6 S) were calculated from the atomic coordinate of AtFN3K using HYDROPRO (*75*). Data fit and c(s) distribution plots were generated using GUSSI (*76*).

### Diamide treatment of HEK293T cells

HEK293T cells were cultured in Dulbecco’s Modified Eagle Media (DMEM) containing 10% Fetal Bovine Serum (FBS) on a 6 cm plate (total 6 plates) and allowed to grow overnight. Cells were transfected with 10 μg of Flag-tagged HsFN3K (EX-W1392-M46, GeneCopoeia) using Calcium Phosphate Transfection protocol (43). Cells were allowed to grow for 48h. After 48h cells were treated with indicated concentrations of Diamide (Sigma) for 2 h. Cells were lysed in buffer containing 50 mM Tris.HCl, pH 7.5, 150 mM NaCl, 10% glycerol, 1% TX-100 and 1x protease inhibitor cocktail (EMD-Millipore). Total Cell Lysate (TCL) was spun at 15,000 rpm for 10 min in a refrigerated centrifuge. Proteins were resolved on a 12% SDS-PAGE gel, transferred on PVDF membrane and detected by Western Blotting using anti-Flag antibody (Cell Signaling Technology).

### CRISPR knockout (KO) cell line and Metabolomics

HepG2 cells were transfected with FN3K Double Nickase Plasmid (sc-412985-NIC) (Santa Cruz Biotechnology) and selected with puromycin according to manufacturer’s protocol. FN3K deletion was confirmed by Western Blotting using an anti-FN3K antibody (Invitrogen) (fig. S7). Stable cell line stocks were made from single cell colony. Cell cultures for both cell lines were grown simultaneously with identical media components. Upon achieving ∼80% confluence in 10 cm culture dish, culture media was removed, and cell monolayer was washed with phosphate buffered saline. Adherent cells were harvested in ice-cold 80% methanol extraction solvent and flash-frozen in liquid nitrogen. Aqueous metabolites were extracted by vortexing/lysing cell pellets in the extraction solvent and collecting the supernatant. Approximately 10% of the supernatant was taken from each sample to form an internal pooled sample. The solvent was then evaporated to produce dried extracts using a CentriVap Benchtop Vacuum Concentrator (Labconco, Kanas City, MO, USA). Extracts were reconstituted in a deuterium oxide (Cambridge Isotope Laboratories) 100 mM phosphate buffer (pH 7.4) with 1/3 mM DSS (Cambridge Isotope Laboratories), which was referenced to 0.0 ppm. ^1^H NMR spectra were acquired on all samples using noesypr1d pulse sequence on an 800 MHz Bruker Avance III HD spectrometer equipped with a SampleJet sample changer (Bruker BioSpin). ^1^H-^13^C heteronuclear single quantum correlation (^1^H-^13^C HSQC) and ^1^H-^1^H total correlation spectroscopy (^1^H-^1^H TOCSY) spectra were acquired on the internal pooled sample for metabolite annotation. Two-dimensional spectra were processed with NMRPipe (*77*), and uploaded to the Complex Mixture Analysis by NMR webserver (COLMARm) (*78*). Metabolite annotations were assigned a confidence score from 1-5 as previously described (*79*) (table S2). ^1^H NMR spectra were phased, and baseline corrected using TopSpin software (Bruker BioSpin) and then further processed and analyzed with an in-house MATLAB toolbox (*80*). After referencing to DSS, end removal, solvent region removal and alignment using the Peak Alignment by Fast Fourier Transform (PAFFT) algorithm (*81*), spectra were normalized using the Probabilistic quotient normalization (PQN) algorithm (*82*). Spectral features that were not significantly overlapped and assigned to annotated metabolites were integrated to get the relative quantification of metabolites for all samples. Annotated metabolites were then tested for significance between cell lines via two-tailed T-test, and p-values were corrected for false discovery rate using the Benjamini-Hochberg method (*83*) (table S2). All spectral data, sample preparation and processing details will be made available on The Metabolomics Workbench (https://metabolomicsworkbench.org).

### Construction of plasmids for transient transformations of N. benthamiana

For constructs used in the transient assays, full-length wild-type or mutant-coding sequences (CDS) of AtFN3K were cloned into pENTR/D-TOPO Gateway entry vector (Invitrogen) following the manufacturer’s protocol. The coding regions were recombined into binary destination expression vector: pSITE-2CA N-terminal green fluorescent protein (GFP) fusion.

*Agrobacterium tumefaciens* strain C58C1 was used for the transient transformation. Colonies carrying the binary plasmids were grown at 28 °C on plates with Lysogeny broth (LB) medium that contained 50 µg/ml gentamycin and 25 µg/ml rifampicin for selection of the strain, and 100 µg/ml spectinomycin for selection of the binary vector. For agroinfiltration, single colonies were grown overnight in 3 ml LB (28°C, 220 rpm). 50 μl of the agro suspension was added to 5 ml LB and the culture was grown overnight. The agro was pelleted by centrifuging at 4,000 rpm for 20 min or 5,0000 rpm for 15 min and the suspension was adjusted to an OD_600_ of 0.2-0.3 in infiltration buffer containing 10 mM MgCl_2_, 10 mM MES, pH 5.7 and 150 mM acetosyringone, pH 5.6, and incubated at room temperature for 3 hours before infiltration. To enhance transient expression of the fusion proteins, the viral suppressor of gene silencing p19 protein was co-expressed. For co-infiltration, equal volumes of cultures were mixed and infiltrated into *N. benthamiana* leaves through the abaxial surface using a 1 ml needleless syringe (Becton, Dickinson, and Company). Plants were then kept in a growth room at 24/22 °C with a 16/8-hour light/dark photoperiod for 48-72 hours.

### Subcellular Localization epifluorescence and confocal microscopy

*N. benthamiana* leaf samples 70 to 90 hours post-infiltration (approximately 0.25 cm^2^ from the infiltrated area) were mounted in water and viewed directly with a Zeiss LSM 880 confocal scanning microscope using an oil immersion objective 40× Plan-Apochromat 1.4NA (numerical aperture of 1.4). Fluorescence was excited using 488 nm light for GFP. GFP emission fluorescence was selectively detected at 490-540 nm using the Zen 2.3 SP1 software. Each experiment was repeated three times.

### Synthesis of ribulose-N-α-Ac-lysine

A solution of *N*-α-acetyl-L-lysine (0.5 g, 2.66 mmol) and D-ribose (1.6 g, 10.64 mmol) in methanol (100 mL) was stirred at 50°C for 4 hours under argon gas. The solvent was evaporated under vacuum and the residue obtained was purified over silica gel column by flash chromatography (R_f_ = 0.14, EtOAc/MeOH/H_2_O, 5/3/2, v/v/v). The product obtained was further purified over a column of Dowex® 50 x 8 H^+^ resin (0.6 x 5 cm) cation exchange resin using 150 mM NaCl as an eluent and desalted over a P-2 column using H_2_O as an eluent. The desired fractions were combined, concentrated and lyophilized to afford the product as white solid (0.55 g, 64.7%); LRMS (ESI): calculated for C13H25N2O7 [M+H]^+^ 321.16, found 321.09. Synthesized ribulose-*N*-α-Ac-lysine was characterized using NMR and ESI (fig. S10) and used as substrate in the PK/LDH assays.

### Statistical analysis

ΔG P-value reported for solvation free energy was calculated by the PISA server (*33*). In brief, it estimates the statistical significance of observing lower than observed ΔG values when the interface atoms are picked randomly from the protein surface with the equivalent interface area. ΔG P-values < 0.5 are statistically significant (*84*).

Statistically significance of metabolite difference between WT and KO was determined using two-tailed T-test. The P-values were then corrected for false discovery rate using the Benjamini-Hochberg method (table S2).

## Acknowledgments

We thank the staff at the Southeast Regional Collaborative Access Team (SER-CAT) at the Advanced Photon Source. We also thank Prof. Emile Van Shaftingen (Université Catholique of Louvain, Brussels, Belgium) for the AtFN3K WT pET-15b construct. **Funding:** This work was initially supported by National Science Foundation (NSF) funds to Kannan lab (MCB-1149106) and more recently by the National Institute of Health (NIH) (R01GM114409) Pump-priming grant from the University of Georgia and University of Liverpool. Additional support for DPB and PAE was received from a Royal Society Research Grant and North West Cancer Research (CR1208). **Author contributions**: N. Kannan conceived and designed the project. S.S. and S.K. performed the experiments. C.E.S.-R. contributed to mutational analysis. D.P.B. and P.A.E. contributed reagents and analyzed redox data. N.R.K., R.K., and Z.A.W. contributed to X-ray crystallography and structural analysis. H.W.K. collected and analyzed sedimentation velocity data. C.P. and A.S.E. contributed to the NMR studies. M.C. and A.S.E. contributed to the metabolomics studies. N. Keyhaninejad and E.V.K. contributed to localization studies on AtFN3K. P.C. and G.J.B. contributed to the synthesis and purification of ribulose lysine. N. Kannan and S.S. wrote the manuscript with contributions from all authors. **Competing interests**: The authors declare that they have no competing interests. **Data and materials availability**: Structural coordinate and structure factors are made available in the Protein Data Bank (PDB code 6OID). All other data needed to evaluate the conclusions in the paper are present in the paper or the Supplementary Materials.

## Supplementary Materials

**Fig. S1.**
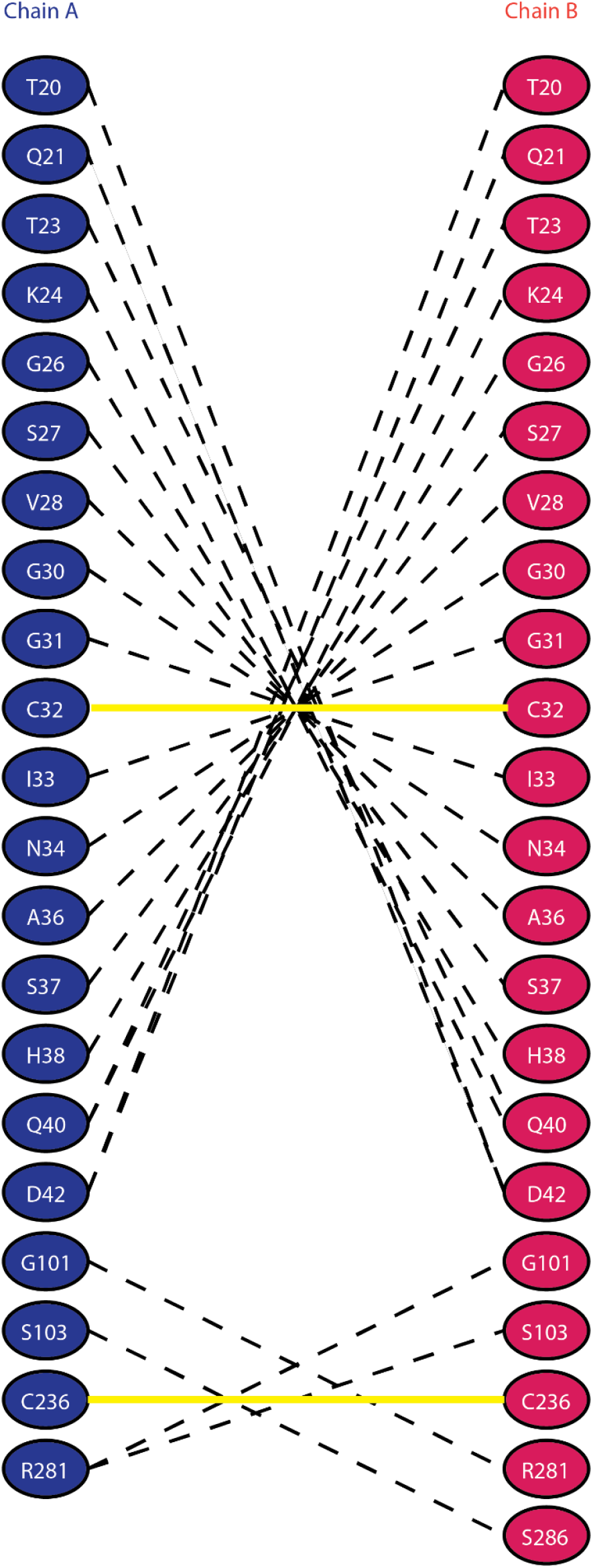
Hydrogen bonds and disulfide bridges between chain A and chain B in WT AtFN3K. The hydrogen bonds between the chains are shown as black dashed lines and disulfide bridges between residues are shown as solid yellow lines. The hydrogen bonds were detected using PISA.

**Fig. S2.**
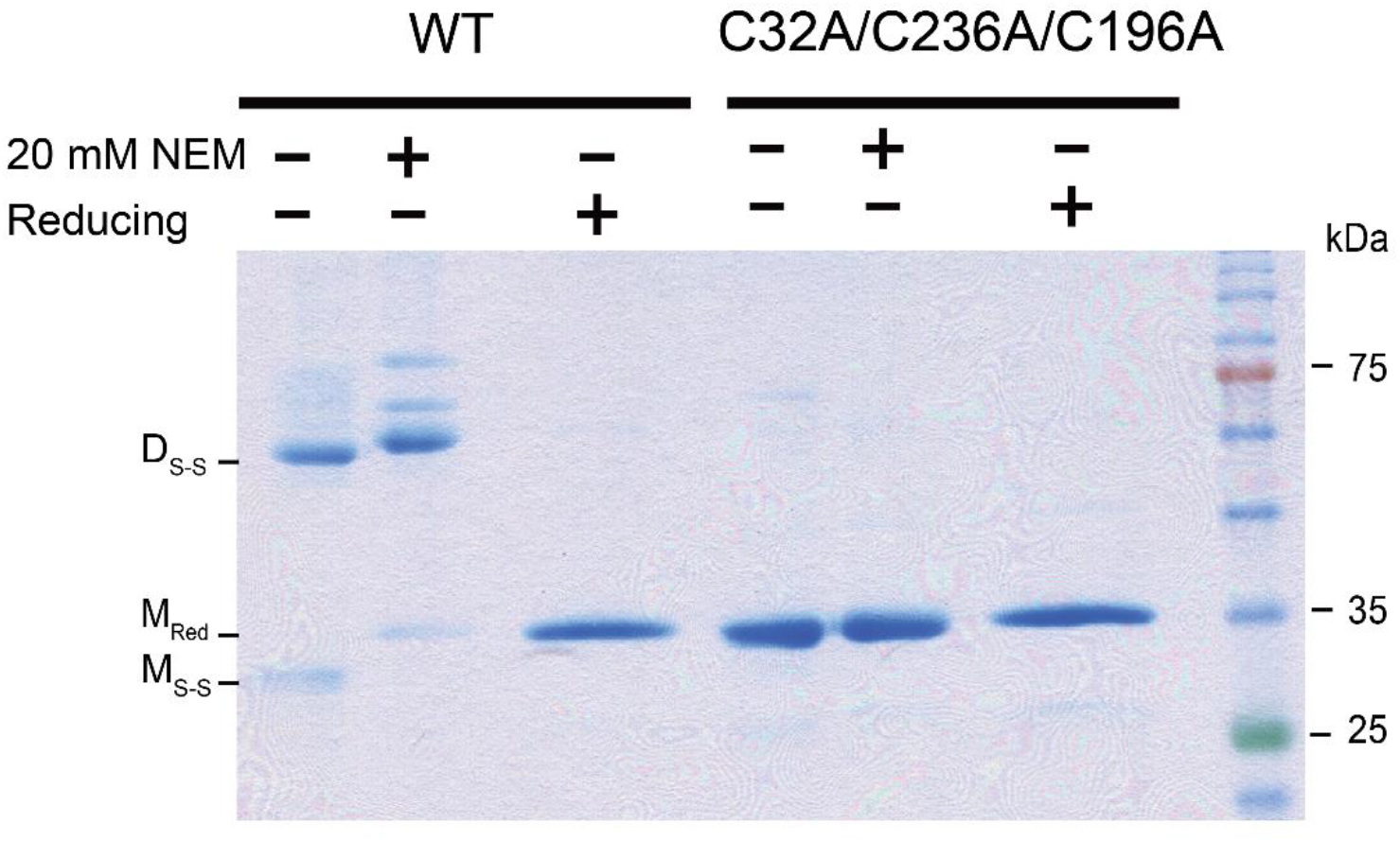
Oxidized monomeric species are an artifact of the SDS-PAGE gel. SDS-PAGE of AtFN3K WT and triple cysteine mutant (C32A/C236A/C196A) dimer species in the presence of N-Ethylmaleimide (NEM) or reducing sample buffer containing 2.8 M 2-Mercaptoethanol (βME). D_S-S_: Disulfide linked dimer, M_Red_: Monomer reduced, M_S-S_: Monomer with intramolecular disulfide. Blot is representative of 2 experiments.

**Fig. S3.**
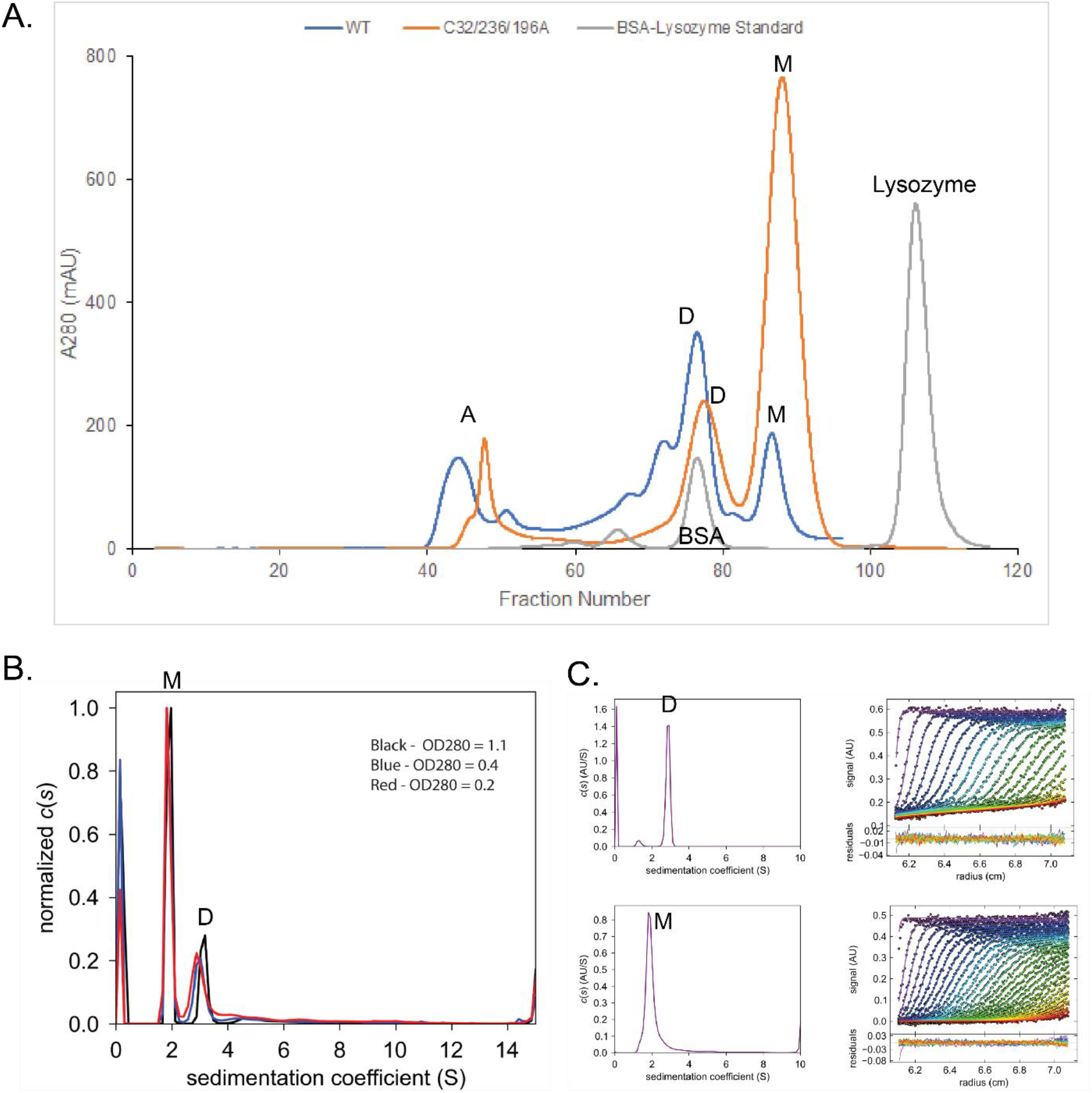
WT and triple-cysteine-mutant (C32A/C236A/C196A) AtFN3K exist as two distinct species. **(A)** Size exclusion chromatography (SEC) of AtFN3K WT (blue) and the mutant (orange) overlaid with the SEC of bovine serum albumin (BSA)-lysozyme mix (gray). AtFN3K WT and triple-cysteine-mutants were purified without co-expression with GroEL chaperone unlike in Fig 6. Inset “A”, aggregates; “D”, dimer; “M”, monomer. **(B)** Sedimentation velocity analysis of the triple cysteine mutant pre-SEC modeled as continuous c(s) distribution. Three protein concentrations were used. Purification and “D and “M” as described/defined in (A). **(C)** Sedimentation velocity analysis of the triple cysteine mutant modeled as continuous c(s) distribution, in which dimer and monomer fractions were isolated using SEC. Purification, D and M as defined in (A). Data in (A to C) are each representative of 2 experiments.

**Fig. S4.**
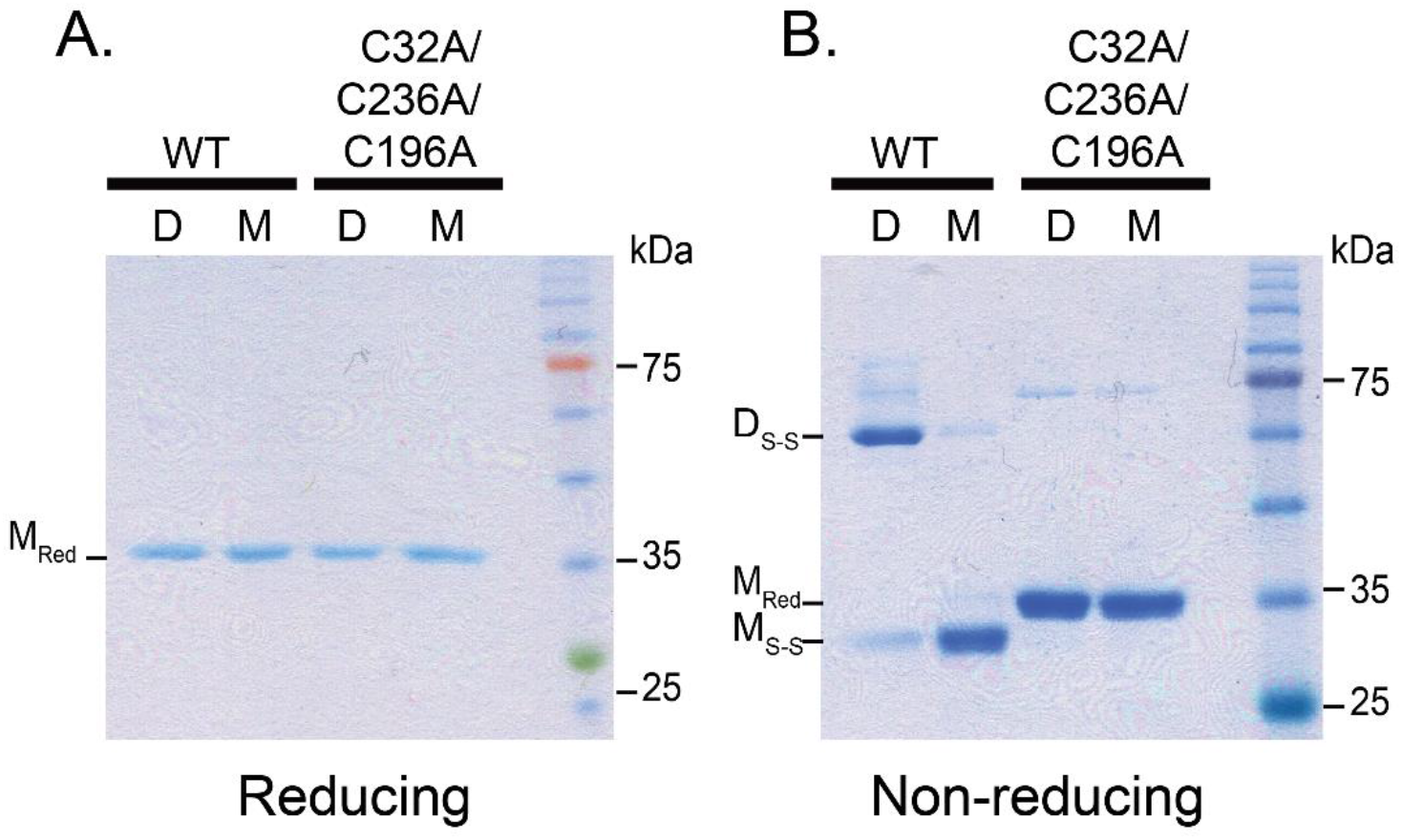
Reducing and non-reducing SDS-PAGE of dimer and monomer fractions of WT and triple-cysteine-mutant AtFN3K. **(A and B)** 3 µg of protein was mixed with either reducing or non-reducing 4X sample buffer and then subjected to SDS-PAGE. D: Dimer Fraction, M: Monomer Fraction, D_S-S_: Disulfide linked dimer; M_Red_: Monomer Reduced; M_S-S_: Monomer with intramolecular disulfide. Approximated 3 µg of protein was loaded. The gel was stained with Coomassie and de-stained with water. Blot is representative of 2 experiments.

**Fig. S5.**
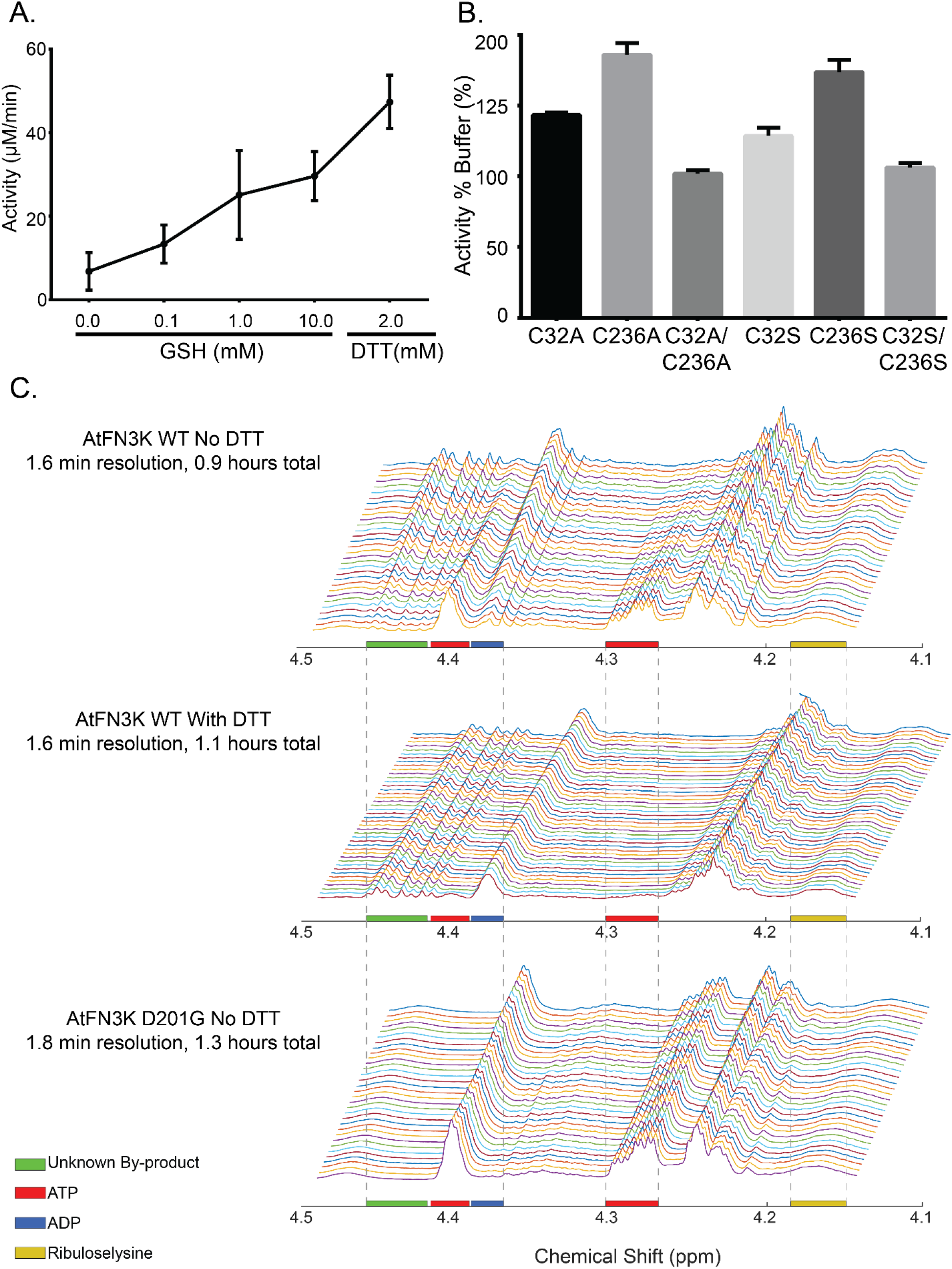
Effects of thiol reagents on activity of WT and cysteine-mutant AtFN3K. **(A)** PK/LDH assay of AtFN3K WT dimer species in the presence of varying concentrations of reduced glutathione (GSH) and 2 mM Dithiothreitol (DTT). **(B)** PK/LDH assay on AtFN3K cysteine mutants in the presence and absence of 2 mM DTT. Activity % Buffer is calculated by dividing activity of the enzyme in the presence of 2 mM DTT by the activity in buffer alone and converting to percentage. Assays were performed with purified protein without SEC. **(C)** NMR spectrum showing the formation of products (ADP and unknown by-product) overtime for AtFN3K WT in the presence and absence of DTT. Kinase-deficient (“dead” mutant D201G) was used as negative control. Assays were performed with purified protein without SEC. Data in (A and B) are means ± SE are each representative, of 3 independent experiments. Data in (C) is representative of 2 independent experiments.

**Fig. S6.**
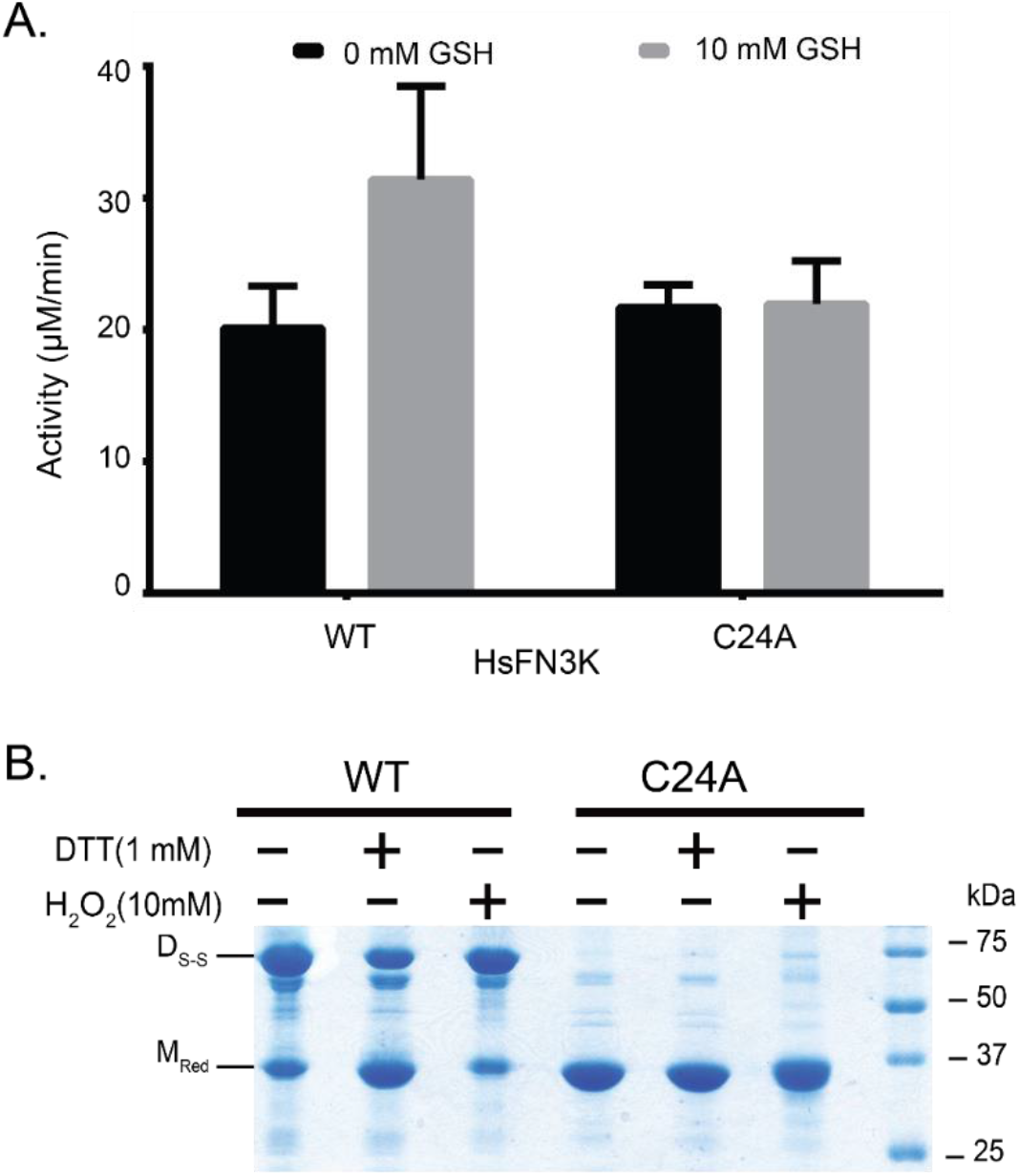
P-loop cysteine (Cys^24^) is critical for the formation of disulfide-linked dimer in HsFN3K. **(A)** PK/LDH assay of HsFN3K WT and C24A in presence and absence of reduced glutathione (GSH). **(B)** Non-reducing SDS-PAGE of HsFN3K WT and C24A mutant. 15 μg of the recombinant protein was incubated with either water, 1 mM DTT or 10 mM H_2_O_2_ for half an hour before adding 4X Sample Buffer. The bands representing disulfide linked dimeric and monomeric species are labelled as D_S-S_ and M_Red_ respectively. The gels were stained with Coomassie Brilliant Blue. The molecular marker is shown on the right. Blots are representative of 3 experiments.

**Fig. S7.**
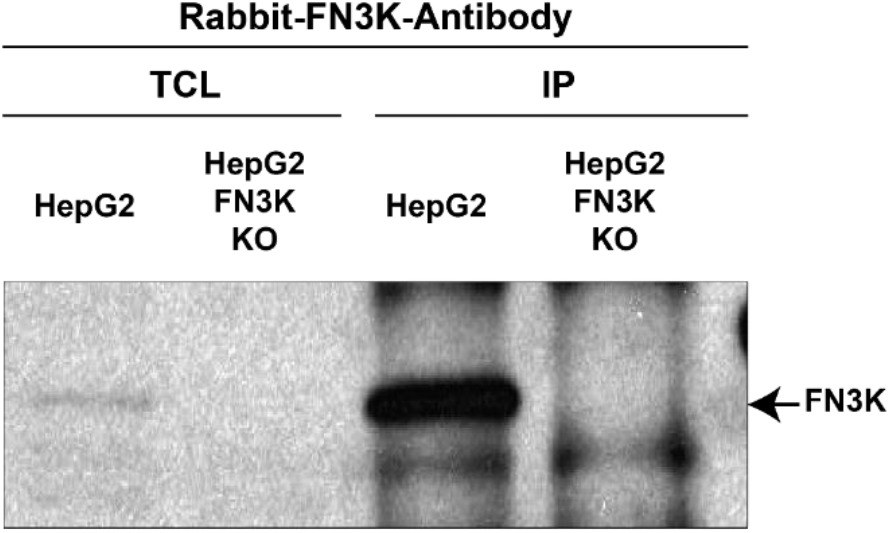
Western blot of HsFN3K KO in HepG2 cells. HepG2 (WT/KO) cells were lysed in lysis buffer (50mM Tris HCl pH 7.4, 100mM NaCl, 10% Glycerol, 1mM EDTA, 1% TritonX-100). Total cell lysate (TCL) was subjected to immunoprecipitation (IP) with anti-FN3K antibody. Proteins were resolved on 12% SDS PAGE gel and immunoblotted with Rabbit-FN3K antibody. Blot is representative of 3 experiments.

**Fig. S8.**
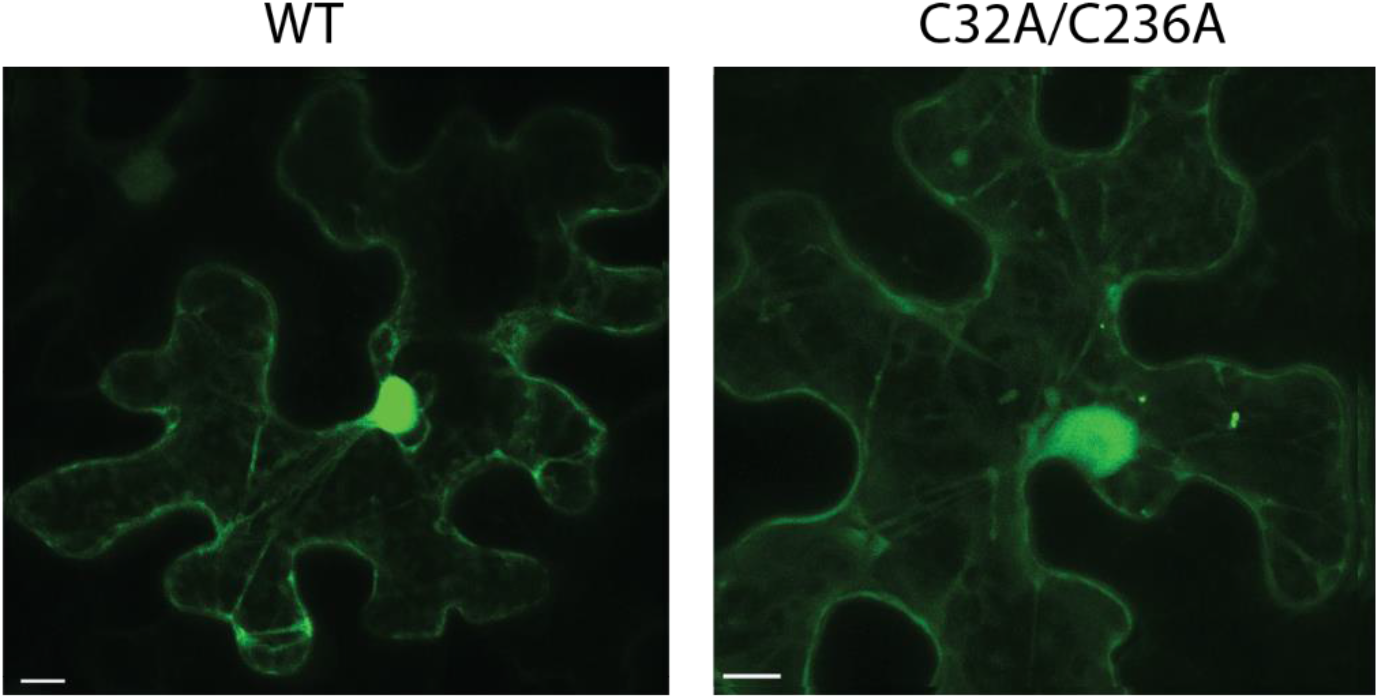
WT and C32A/C236A AtFN3K localizes to the nucleus. Subcellular localization of AtFN3K WT and C32A/C236A respectively, in *N. benthamiana* leaf epidermal cells. Scale bar:10 μm. Images are representative of 3 experiments.

**Fig. S9.**
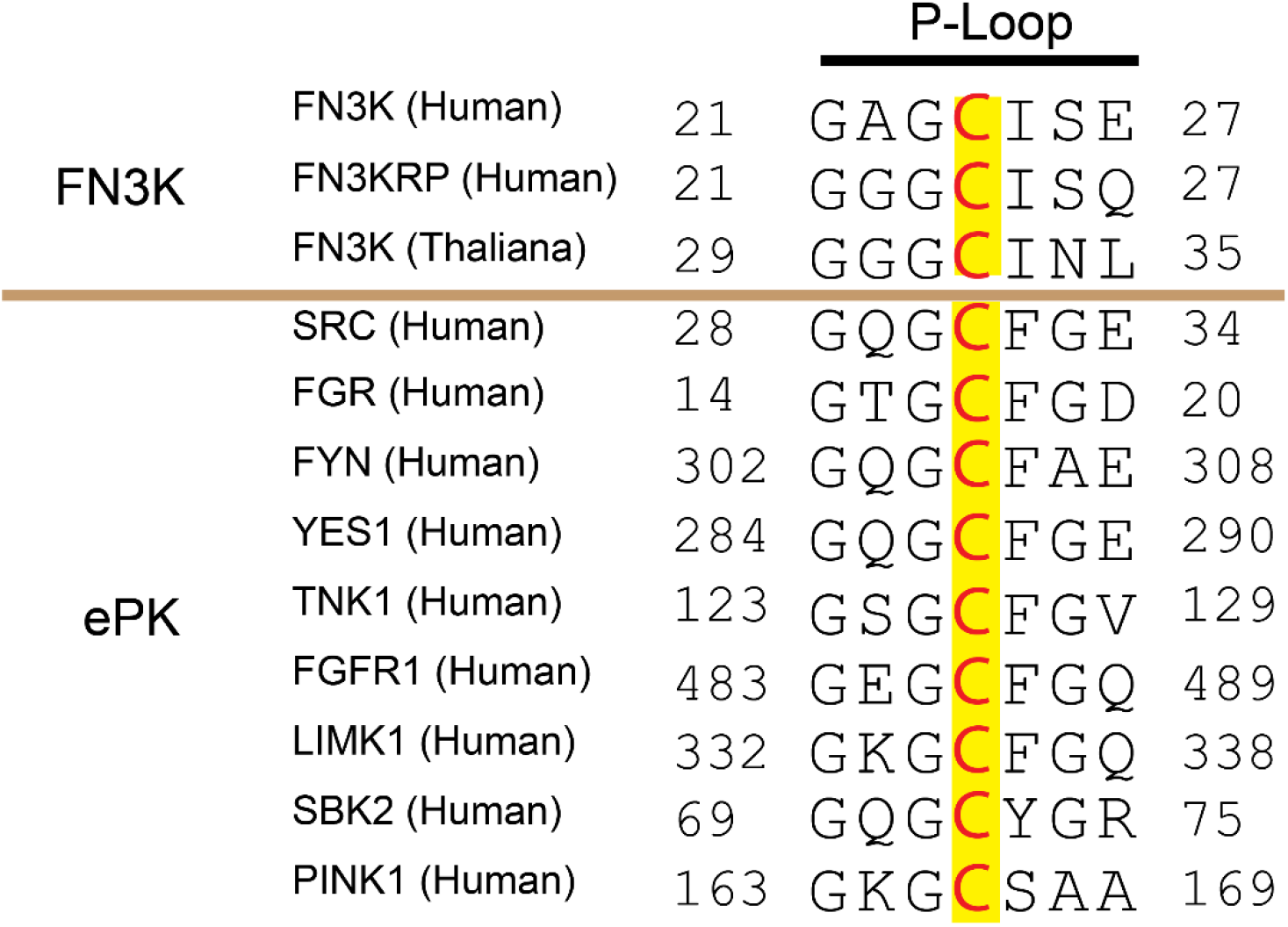
Multiple sequence alignment showing the conservation of P-loop cysteine in selected human eukaryotic Protein Kinases (ePKs) and FN3Ks. The human kinases were identified using previously curated alignment of ePKs and small molecule kinases.

**Fig. S10.**
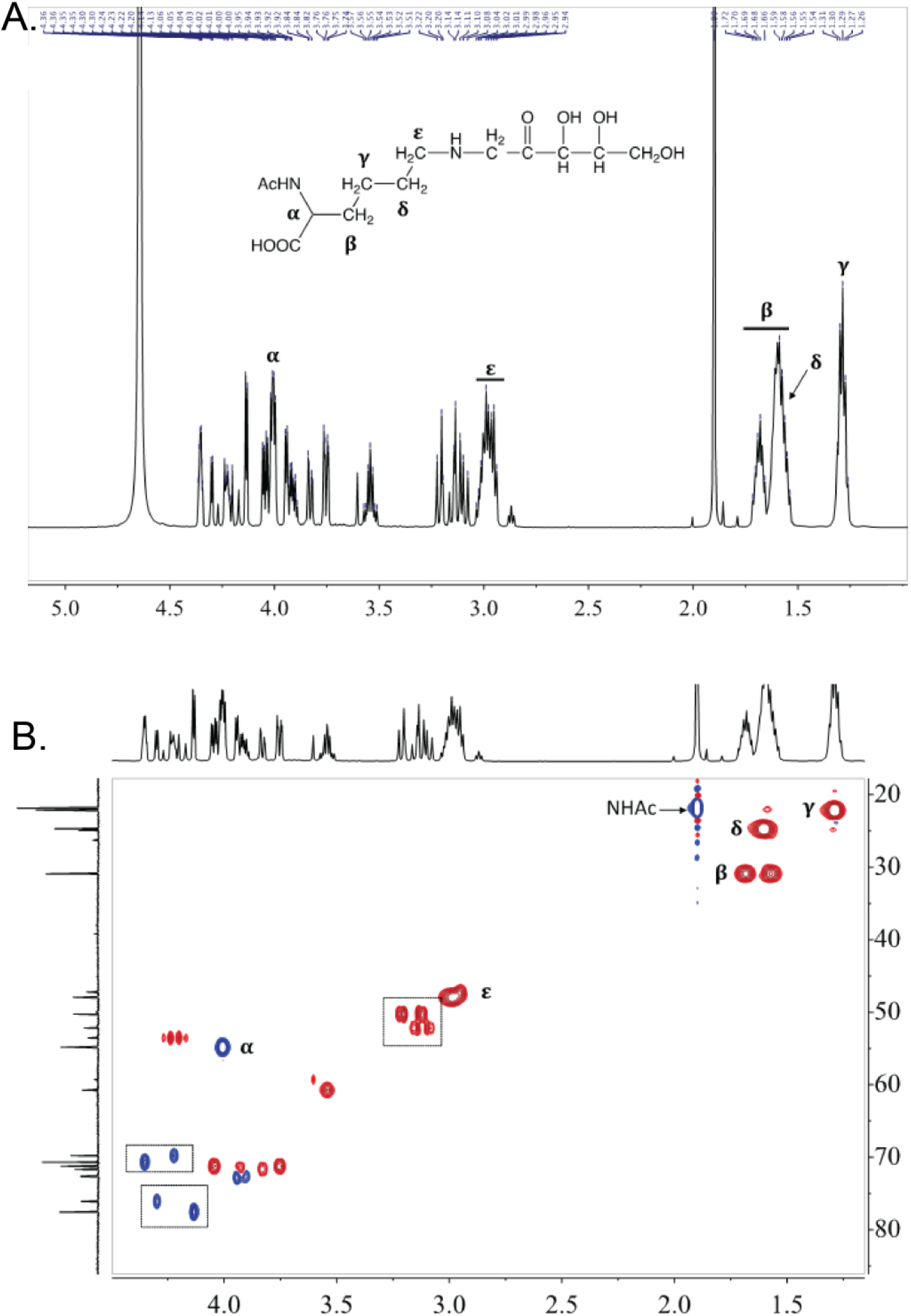
Spectral data of ribulose-N-α-Ac-lysine. **(A)** 1D proton nuclear magnetic resonance spectra (^1^H NMR, 600 MHz, D_2_O) confirms purity and identity of ribose-lysine adduct; open-chain structure in keto form of Amadori rearrangement product is shown; characteristic protons of lysine backbone are labelled; **(B)** 2D heteronuclear single quantum coherence spectra (HSQC, 151 MHz, D_2_O) shows correlation between ^1^H and ^13^C; each peak represents a bonded C-H pair, red peaks corresponds to secondary carbon (CH_2_) and blue peaks corresponds to primary/tertiary carbon (CH_3_/CH); characteristic carbons of lysine backbone are labelled; dashed boxes over sugar carbons highlights that adduct is present in multiple forms.

**Fig. S11.**
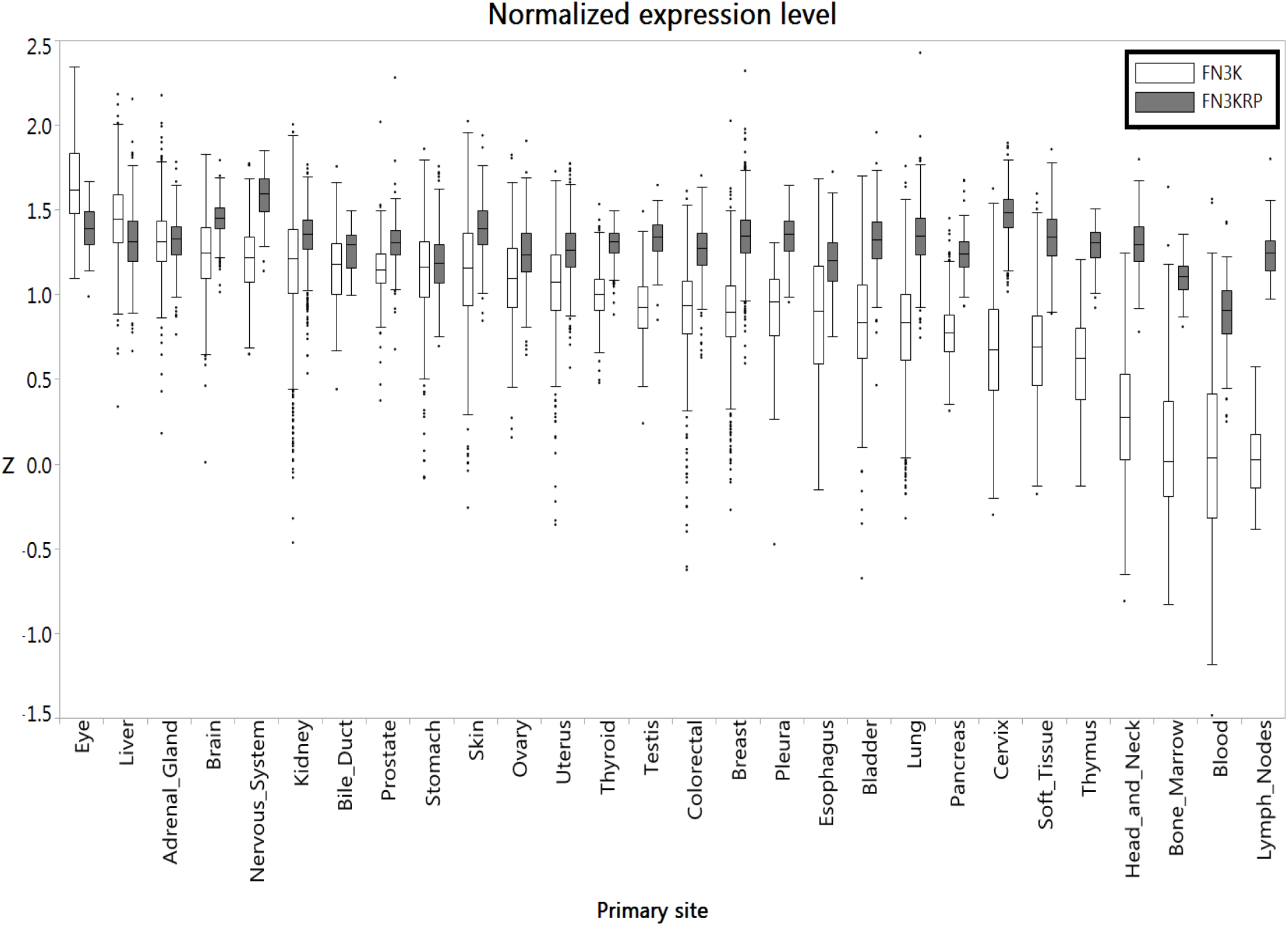
FN3K and FN3KRP expression levels in human tumors. Boxplots show the Z score distributions (Y-axis) of HsFN3K (white boxes) and HsFN3KRP (gray boxes) in different primary tumor sites in descending order of the median Z score. Outliers are marked as dots.

**Table S1.** Data collection and refinement statistics of WT AtFN3K. ^a^Values in parentheses are for highest-resolution shell. ^b^R_meas_ is the redundancy independent merging R-factor of Diederichs and Karplus (*88*). ^c^CC_1/2_ is the percentage of correlation between intensities from random half-data sets (*89*).

| **Data Collection** |  |
| --- | --- |
| Space Group | C 2 2 21 |
| Unit cell dimensions (a, b, c, (Å) $\alpha$, $\beta$, $\gamma$) | 158.86 163.64 70.72<br>90.00 90.00 90.00 |
| Total reflections | 551252 (39245) |
| Unique reflections | 38000 (2683) |
| Redundancy | 14.5 (14.6) |
| Completeness (%) | 1.00 (0.97) |
| I/$\sigma$(I) | 14.3 (1.44) |
| R-meas<sup>b</sup> (%) | 0.157 (2.50) |
| CC<sub>1/2</sub><sup>c</sup> (%) | 0.999 (0.519) |
| **Refinement** |  |
| Resolution (Å) | 53.50 - 2.37 |
| R<sub>work</sub> / R<sub>free</sub> | 0.175 / 0.204 |
| No. atoms<br>Protein/Ligand/Water | 4583/154/89 |
| B-factors (Å<sup>2</sup>)<br>Protein/Ligand/Water | 64.1/92.4/56.3 |
| **Stereochemical Ideality** |  |
| Bond lengths (Å<sup>2</sup>) | 0.009 |
| Bond angles (°) | 1.07 |
| Ramachandran favored (%) | 97 |
| Ramachandran allowed (%) | 3 |
| Ramachandran outliers (%) | 0 |

**Table S2.**
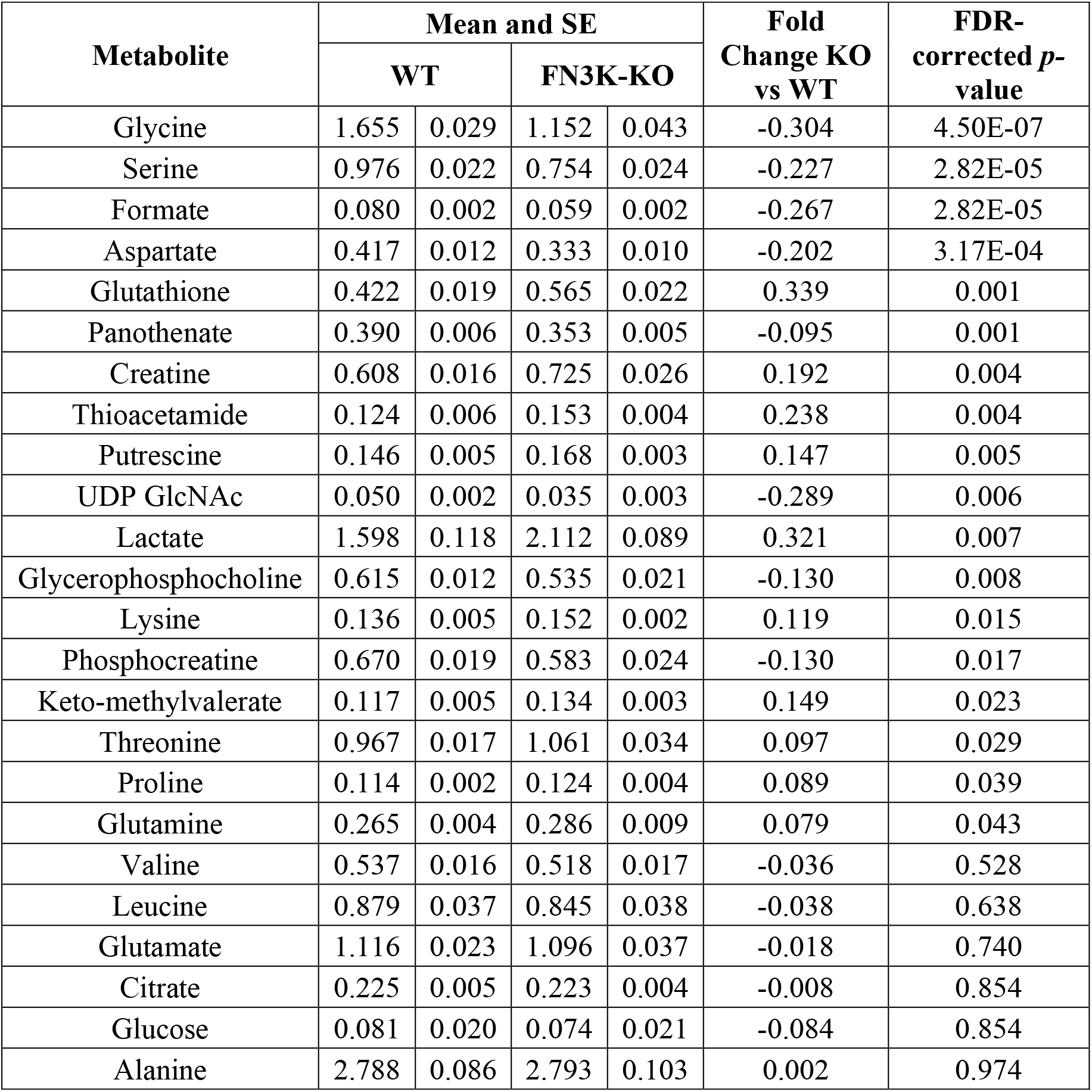
Metabolites identified in extracts of WT HepG2 and FN3K-KO cells. ^a^ Confidence scores are defined as follows: 1. putatively characterized compound classes or annotated compounds, 2. matched to literature and/or 1D database, 3. matched to 2D ^13^C-^1^H HSQC, 4. matched to HSQC and validated by second 2D dataset (^1^H-^1^H TOCSY), and 5. validated by spiking the authentic compound into sample; ^b^ Mean and standard error (SE) values are reported in relative arbitrary units, calculated after spectral normalization, n=10 and 9 for WT and KO, respectively; ^c^ FC: fold change, calculated as ratio of KO mean to WT mean; ^d^ UDP GlcNAc: uridine diphosphate N-acetyl-glucosamine.

